# Respiratory viruses activate autophagy via the IFN-STAT1/STAT5B-SOCS1 axis

**DOI:** 10.1101/2025.10.28.685013

**Authors:** Victoria Hunszinger, Helen Dürr, Zoé Engels, Helene Hoenigsperger, Lennart Koepke, Susanne Klute, Jana-Romana Fischer, Birgit Ott, Alexander Graf, Stefan Krebs, Helmut Blum, Maximilian Hirschenberger, Frank Kirchhoff, Konstantin MJ Sparrer

## Abstract

Autophagy is an ancient catabolic process that has emerged as part of the innate immunity. Upon infection, autophagy is activated but key factors responsible remained unclear. Here, we show that interferon (IFN) released during viral infections subsequently activates autophagy via STAT1/5B-mediated upregulation of SOCS1. Our data shows that scavenging of IFNs diminishes autophagy induced by several respiratory viruses. All types of IFN (I, II and III) mediated robust autophagic flux activation in both cell lines and primary human lung fibroblasts in a JAK1-3 dependent manner. Depletion or pharmacological inhibition of individual STAT transcription factors demonstrated that both STAT1 and STAT5B are required for IFN-induced autophagy. Upon IFN stimulation STAT1 and STAT5B associate and translocate to the nucleus. Transcriptome analyses revealed that inhibition of STAT5 only reduces expression of a subset of IFN-stimulated genes, whereas most known anti-viral IFN-stimulated genes remain induced to high levels. Among the STAT5 dependent genes was Suppressor of Cytokine Signaling 1 (SOCS1). Overexpression of SOCS1 stimulated autophagy, whereas its depletion impaired IFN-induced autophagy. Successful viruses like measles virus (MeV) or respiratory syncytial viruses (RSV) evolved strategies to exploit autophagy to promote their own replication. Uncoupling IFN-mediated ISG defenses from autophagy induction using by STAT5 inhibition reduced virus-induced autophagy, and inhibited efficient replication of autophagy-dependent MeV and RSV. Taken together, our data show that IFN promotes autophagy via STAT1/STAT5B-SOCS1 in viral infections and reveal that targeting of this axis allows inhibition of autophagy-dependent viruses without compromising innate immune defenses.

**Author summary:** Autophagy, like the Interferon (IFN) system, is considered part of the innate immune defenses that are activated upon viral infection. However, some viruses—like measles (MeV) and respiratory syncytial virus (RSV)—have learned to hijack this process to promote their replication. Our study shows that autophagy activation upon viral infections is promoted by the IFN system. Consequently, blocking IFNs reduced virus-induced autophagy. Mechanistic analyses demonstrated that various types of IFN induce expression of SOCS1, via the transcription factors STAT1 and STAT5B in cell lines and primary human lung cells. Pharmacological inhibition or depletion of STAT1, STAT5B or SOCS1 impaired IFN-induced autophagy. Importantly, disrupting STAT5 signaling does not impact IFN-mediated induction of the large majority of ISGs. Thus, selective disruption of IFN-induced autophagy by inhibiting STAT5 restricted replication of MeV and RSV. Our findings reveal an underappreciated mechanistic link between the IFN system and autophagy and suggest that targeting of the STAT5-SOCS1 axis may inhibit autophagy-dependent viruses without compromising IFN-mediated defenses.

## Introduction

Innate immunity is one of the major defense systems against invading pathogens, including respiratory viruses [1,2]. Activation of innate immune pathways such as the interferon (IFN) system or autophagy during viral infection is a highly coordinated process. However, while activation of the IFN system is considered anti-viral against most respiratory viruses, autophagy has been usurped by successful viruses such as Measles Virus (MeV) and Respiratory Syncytial Virus (RSV) to promote their replication.

The innate immune system detects viral infections mainly via pattern recognition receptors (PRR), such as retinoic acid-inducible gene (RIG)-I-like receptors (RLRs), Toll-like receptors (TLRs), or cyclic GMP-AMP synthase (cGAS) that recognize both intracellular and extracellular viral pathogens by binding to pathogen-associated molecular patterns (PAMPs) [2–6]. Activation of PRRs converges in the induction and secretion of IFNs and other pro-inflammatory cytokines. The three types of IFNs (type I, II or III) include the type I IFNs, IFN-α and -β, the sole type II IFN, IFN-γ, and the four type III IFNs, IFN-λ1-4. Of note, while type I IFNs can be secreted by almost all cell types, type II IFN production is mainly limited to lymphocytes, whereas type III IFNs are primarily found at mucosal surfaces [7,8]. The secreted IFNs bind to their respective receptors in both an autocrine and paracrine fashion and mediate activation of a set of Janus kinases (JAK1-3 and Tyk2) [9,10]. The kinases in turn phosphorylate the signal transducer and activator of transcription (STAT) transcription factors. Humans express seven different STAT proteins, namely STAT1, STAT2, STAT3, STAT4, STAT5A, STAT5B and STAT6 [11]. Each of these transcription factors can act as a homodimer or a heterodimer, driving or restricting the expression of a diverse array of genes. For example, type I or III IFN stimulation typically leads to the formation of STAT1-STAT2 heterodimers [12], which eventually activate transcription driven by ISRE (Interferon-Stimulated Response Element) elements. Expression of ISGs, among them many well-known virus restriction factors, such as Mx1 or OAS1, sets the cells in an anti-viral state [13,14].

Macroautophagy (hereafter autophagy) is an evolutionarily conserved and intricately regulated degradation pathway and anti-viral defense mechanism [15–17]. Activation of autophagy is induced by cellular kinases, including ULK1 (unc-51 like autophagy activating kinase 1) and 5’ AMP-activated protein kinase (AMPK), and downregulated by mTOR and Casein kinase II [18,19]. The precise mechanisms by which autophagy is activated during viral infection, however, remain unknown. During autophagy, cytoplasmic cargoes are engulfed by double-membrane vesicles, termed ‘autophagosomes’, which are eventually degraded upon fusion with lysosomes [20]. Processing of LC3B from cytosolic LC3B-I to autophagosome-associated LC3B-II and degradation of the autophagy receptor SQSTM1/p62 are hallmarks of autophagy. Autophagy targets viruses or viral components by dedicated receptors, such as p62 to promote their lysosomal degradation [16,20,21]. Unfortunately, the ancient autophagy system and its components have been hijacked and exploited by viruses during their long co-evolution, effectively making autophagy a pro-viral process for some viruses. For example, interaction of the C protein of MeV with IRGM promotes pro-viral autophagy enhancing production of virions (*20*–*22*). Similarly, RSV exploits the autophagic machinery to form cholesterol-rich lysosomes that promote replication [25]. Targeting pro-viral autophagy may thus be a feasible anti-viral strategy.

Autophagy and the IFN system are highly interconnected. For example, activation of the cGAS-STING cascade that mediates sensing of virus-derived cytosolic DNA, induces a strong autophagic response [26]. In line with this, RNA sensing, e.g. by the sensor RIG-I, was also recently suggested to promote autophagy activation [27]. Furthermore, activation of Toll-like receptors such as TLR7, was shown to promote MyD88-dependent autophagy induction [28]. In addition, key factors of the type I IFN induction cascade play roles in both autophagy and the IFN system [13,15]. For example, the kinase TBK1, which is crucial for activation of the transcription factor IRF3, was reported to mediate virus-induced autophagy by phosphorylating the autophagy receptor p62 [2,29,30]. *Vice versa*, autophagy was reported to dampen IFN induction by mediating the turnover of signaling components or effector proteins [15]. For example, the DNA sensor cGAS and the RNA sensor RIG-I are degraded by autophagy [31,32]. It was suggested that IFNs itself may also promote autophagy [33,34]. However, the underlying mechanism(s) are currently not understood.

Here, we show that induction of autophagic flux during respiratory viral infection is activated by virus-induced IFNs. Mechanistic analyses revealed that JAK-STAT1/5B-SOCS1 signaling is required to activate autophagy. Notably, we further show that selectively inhibiting IFN-induced autophagy by targeting of STAT5B signaling does not impair IFN-mediated defenses but interferes with replication of autophagy-dependent viruses such as MeV and RSV. Our results establish a connection between IFN signaling and autophagy via the STAT1/5B-SOCS1 axis that contributes to autophagy induction during viral infections. Targeting of this axis inhibits replication of autophagy-dependent viruses.

## Results

### Autophagy induction during viral infection is dependent on IFNs

To understand whether autophagy induction during viral infection is coordinated with the IFN system, we inhibited type I IFN by B18R, a potent scavenger derived from a pox virus [35]. Autophagosome levels in infected HeLa autophagy reporter cells stably expressing GFP-LC3B were analyzed by flow cytometry [36]. To this end, we mildly permeabilized the cells and removed non-membrane-bound GFP-LC3B-I by washing. Quantification of the remaining fluorescence is a proxy for the GFP-LC3B-II-positive autophagosome content [36]. Inhibition of IFN by B18R reduced autophagosome levels in MeV-, RSV- or IAV-infected GFP-LC3B expressing HeLa cells significantly (Fig 1A). This was confirmed by analyzing GFP-LC3B puncta as surrogate for autophagosomes in HeLa reporter cells infected with MeV, RSV and IAV via immunofluorescence (Fig 1B). B18R treatment efficiently reduced the amount of autophagosomes after infection almost to baseline levels (Fig 1C). These data indicate that activation of autophagy during respiratory viral infection is further driven by infection-induced IFNs.

**Fig 1.**
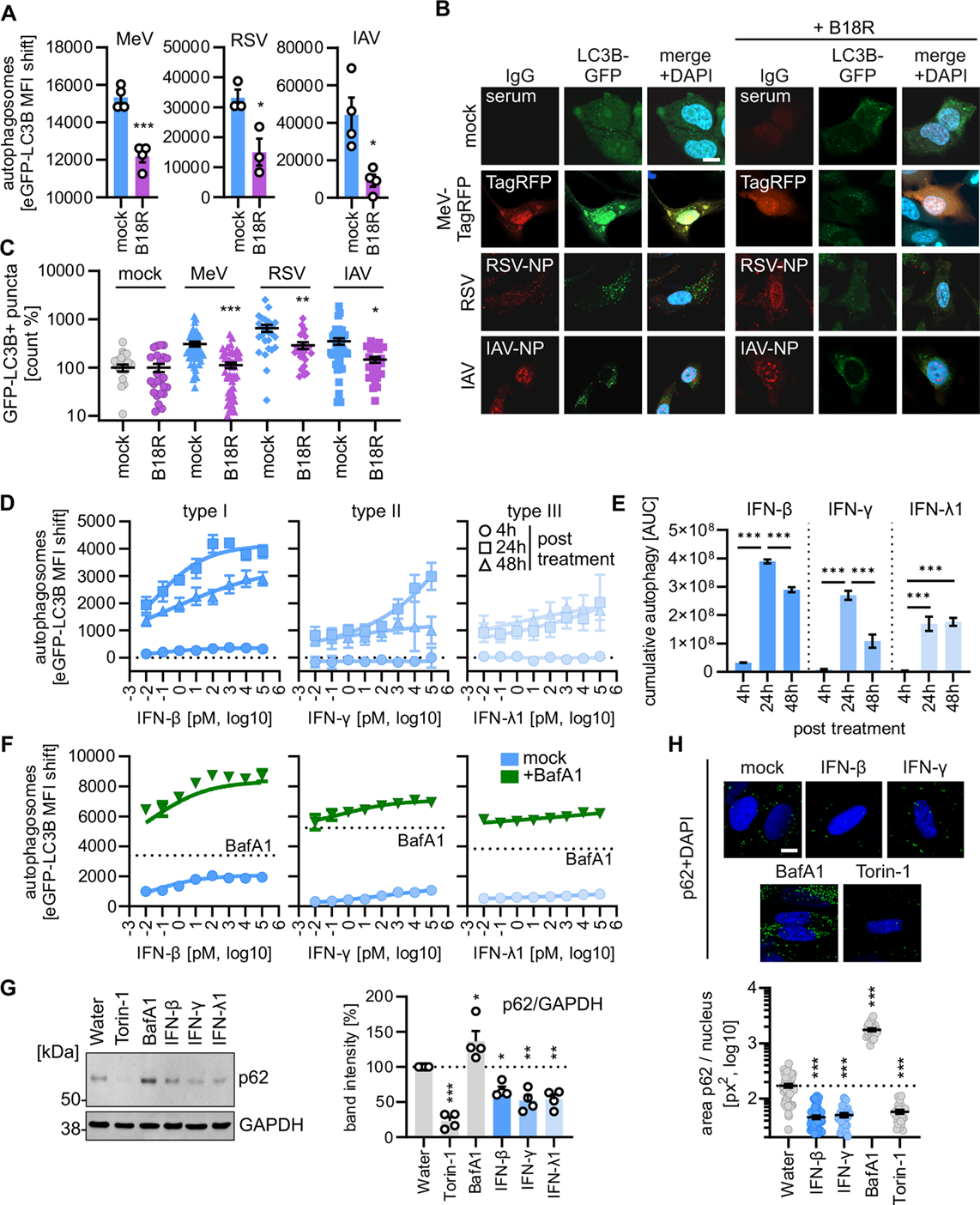
Interferons induce autophagic flux. **A,** Quantification of autophagosome levels by flow cytometry in HeLa autophagy reporter cells (HeLa GL) 24 h after infection with Measles Virus (MeV, MOI 1.25), Respiratory Syncytial Virus (RSV, MOI 1.25), or Influenza A Virus (IAV, MOI 1.25). Treated with 200 ng/ml B18R as indicated. n = 3 – 4 biological replicates ± SEM. Student’s t-test with Welch’s correction. **B,** Representative confocal immunofluorescence images of HeLa autophagy reporter cells (HeLa GL, GFP-LC3B, green) 24 h after infection with Measles Virus expressing TagRFP (MeV-TagRFP, MOI 1), Respiratory Syncytial Virus (RSV, MOI 1), or Influenza A Virus (IAV, MOI 1) treated with 200 ng/ml B18R as indicated. **C,** Quantification of the number of autophagosomes (= eGFP-LC3B positive puncta) per nucleus per image relative to the uninfected controls of (B) n= 25-50 images ± SEM. Brown-Forsythe and Welch ANOVA with Dunnett’s T3 multiple comparisons test. **D,** Quantification of autophagosome levels by flow cytometry in HeLa autophagy reporter cells (HeLa GL) 4, 24 and 48 h after treatment with increasing concentrations of indicated IFNs (0.01 pM – 100 nM). n = 4 ± SD. **E,** Area under the curve analysis showing accumulated autophagy levels over time. Accumulated data of (D). Bars indicate the means of n = 4 ± SEM. Brown-Forsythe and Welch ANOVA with Dunnett’s T3 multiple comparisons test. **F,** Quantification of autophagosome levels by flow cytometry in HeLa autophagy reporter cells (HeLa GL) 24 after treatment with increasing concentrations (0.01 pM – 100 nM) of indicated IFNs in presence of Bafilomycin A1 (625 nM). Dotted line, Bafilomycin A1 only treatment. n = 4 ± SD. **G,** Representative immunoblots of NHLF cells treated for 48h with Torin-1 (1 µM), Bafilomycin A1 (250 nM) or 100 pM of either IFN-β, IFN-γ or IFN-λ1. Stained with α-SQSTM1/p62 and α-GAPDH (left panel). Quantification of SQSTM1/p62 band intensities normalized to GAPDH band intensities of the immunoblots as percentage of the water control. n=4 (biological replicates) ± SEM (right panel). Brown-Forsythe and Welch ANOVA with Dunnett’s T3 multiple comparisons test. **H,** Representative confocal immunofluorescence images of NHLF cells treated with IFN-β or IFN-γ (each 100 pM) for 24 h. BafA1 (625 nM, 4h), Torin-1 (1 µM, 4h) were used as controls. Stained with α-SQSTM1/p62 (green) and DAPI (blue) (scalebar 10 µm, left panels). Quantification of the number of autophagosomes (= SQSTM1/p62 positive puncta) pr nuclei per tile (right panel). n= 25-50 tiles ± SEM. Brown-Forsythe and Welch ANOVA with Dunnett’s T3 multiple comparisons test. *, p<0.05, ** p<0.01, *** p<0.001.

### Various types of IFNs promote autophagic flux

To determine whether the link between autophagy and IFN is dependent on the type of IFN, we treated GFP-LC3B expressing HeLa autophagy reporter cells with selected type I, type II or type III IFNs (type I: IFN-β, IFN-α2, IFN-ε; type II: IFN-γ; type III: IFN-λ1, IFN-λ2/3). Treatment with all tested IFNs increased the number of autophagosomes in HeLa cells in a dose-dependent manner (Figs 1D and S1A). Induction of autophagosomes was apparent 24h post-treatment for all IFNs. IFN-β and IFN-γ peaked at 24 h, whereas IFN-λ1 induced autophagy stayed constant at lower levels between 24h and 48h post-treatment (Figs 1E and S1A). Detailed analyses of the kinetics of autophagosome induction by IFN-β showed a significant increase of autophagosomes already at 8h post-treatment (Fig S1B). To discriminate whether IFN treatment induced *de novo* autophagic flux or whether autophagosomes accumulated due to inhibited or slowed turnover, we blocked autophagosome turnover using saturating concentrations of Bafilomycin A1 (BafA1) [19,20,37]. Bafilomycin A1 is a potent V-ATPase inhibitor which prevents autophagic flux, leading to the accumulation of autophagosomes. Thus, in the presence of saturating concentrations of Bafilomycin A1 any other compound impairing turnover is masked, but *de novo* autophagy induction is still detectable. These assays revealed that IFN-α2, IFN-β, IFN-ε, IFN-γ, IFN-λ1 and IFN-λ3 induce *de novo* autophagic flux in a dose-dependent manner (Figs 1F and S1C). To establish physiological relevance, we treated primary human lung fibroblasts (NHLF) with type I, II or III IFNs. All treatments significantly reduced SQSTM1/p62 levels 48h post treatment, a hallmark of autophagic flux induction (Fig 1G). Torin-1, a well-known mTOR inhibitor and autophagy-inducer was used as a positive control, whereas BafA1 was used as a negative control (Fig 1G). To corroborate this data, endogenous p62 abundance in NHLF cells was assessed using immunofluorescence. In line, treatment with type I or II IFNs significantly reduced the p62 area per nucleus, similar as treatment with Torin-1 (Fig 1H). These experiments demonstrate that all types of IFNs induce autophagic flux in human cell lines and primary cells.

### IFN-induced autophagy is dependent on JAK1-3

After binding to their respective receptors, IFNs activate the kinases JAK1-3 and Tyk2 (Fig 2A) [2]. They, in turn, phosphorylate and activate the STAT transcription factors, which eventually mediate transcriptional induction of ISGs (Fig 2A). To dissect which of these kinases are required for autophagy induction by IFNs, we used inhibitors: TC JL 37 (TC) targets TYK2, Ruxolitinib (Rux) prevents activation of JAK1/2, PF06651660/Ritlecitinib (PF) inhibits JAK3 and CP690550/Tofacitinib (CP) inactivates JAK1-3 (Fig 2A) [11]. To quantify their impact on IFN-induced autophagy, HeLa autophagy reporter cells were pre-treated with the inhibitors at approximately double the concentration of their respective IC_50_ and then incubated with IFNs. These assays revealed that autophagy induction by all types of IFN is inhibited by the JAK kinase inhibitors PF, Rux and CP, whereas the TYK2 inhibitor TC had lesser effects (Figs 2 B-D and S2 A-C). This indicates that mainly JAK kinases (JAK 1-3) mediate IFN-induced autophagy, whereas TYK2 plays a minor role.

**Fig 2.**
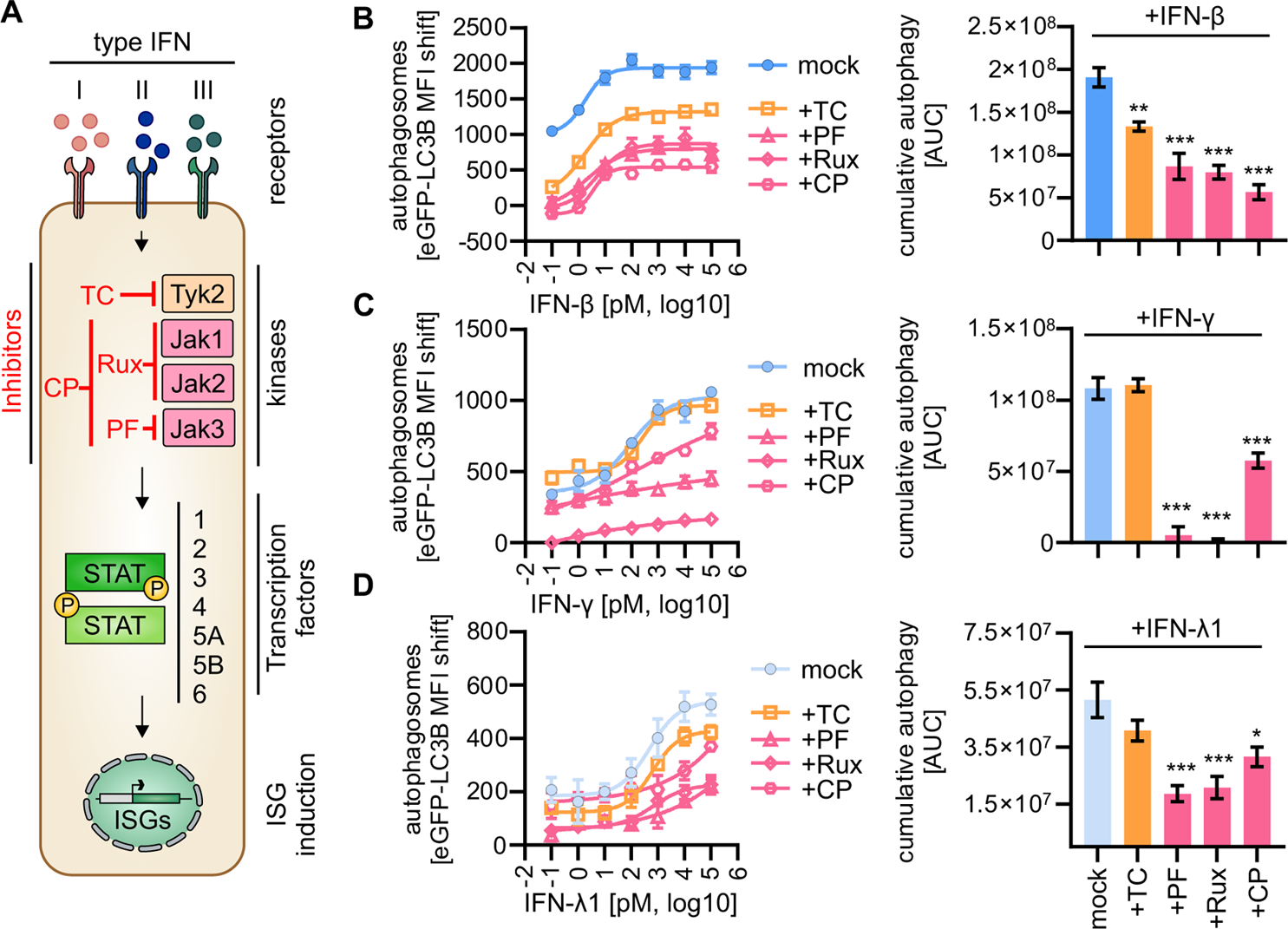
Inhibition of IFN induced autophagy by JAK Inhibitors. **A,** Schematic depiction of the JAK-STAT signaling cascade. First level shows the Type I (red), II (blue) and III (green) IFNs with respective receptors. Second level depicts the receptor-associated kinases (right) Tyrosine kinase 2 (Tyk2), Janus kinase 1 (JAK1), Janus kinase 2 (JAK2) and Janus kinase 3 (JAK) with the respective inhibitors (left) for each kinase. TC JL 37 (TC) inhibits Tyk2, Ruxolitinib (Rux, international nonproprietary name) inhibits JAK1 and JAK2, PF06551600 (PF, international nonproprietary name Ritlecitinib) inhibits JAK3 and CP-690550 (CP, international nonproprietary name Tofacitinib) inhibits JAK1, JAK2 and JAK3. Third level shows exemplary Signal Transducer and Activator of Transcription (STAT) dimer with STAT proteins (STAT1 to STAT6) listed. Fourth level depicts schematic induction of interferon-stimulated gene (ISG) expression in the nucleus. **B-D**, Quantification of autophagosome levels by flow cytometry in HeLa autophagy reporter cells (HeLa GL) 24 h after treatment with increasing concentrations of IFN-β (B), IFN-γ (C) or IFN-λ1 (D), in presence or absence of TC (100 nM), PF (100 µM), Ruxolitinib (Rux, 100 nM), or CP (100 nM), n = 4 ± SEM (left panels). Area under the curve (AUC) analysis (right panels) of the cumulative autophagy induction. Brown-Forsythe and Welch ANOVA with Dunnett’s T3 multiple comparisons test. *, p<0.05, ** p<0.01, *** p<0.001.

### STAT1 and STAT5B are required for IFN-induced autophagy

To investigate which of the seven human STAT proteins are required for IFN-induced autophagy, we constructed a library expressing Cas9 and at least two gRNAs against each STAT protein. Autophagosome induction by IFN-β was abrogated upon expression of gRNAs targeting STAT1 and STAT5B (Fig 3A). Targeting of all other STAT proteins did not have a significant effect. In line, depletion of STAT1 and STAT5B, but not STAT3, by siRNA reduced the autophagosome levels induced by IFN-β treatment in Hela autophagy reporter cells (Fig 3B). Knockdown efficiency was above 90% (Fig S3A). To corroborate this data, we used small compounds that target individual STATs. STAT1 was inhibited by treatment with Fludarabine (STAT1i), STAT5 by CAS 285986-31-4 (STAT5i), STAT3 by 5,15-DPP (STAT3i), and STAT6 by AS1517499 (STAT6i). Treatment of HeLa autophagy reporter cells with STAT1i and STAT5i, but not STAT3i or STAT6i, decreased IFN-β-induced autophagy (Figs 3C, D). Autophagic flux stimulation by IFN-γ and IFN-λ1 was also significantly reduced by STAT5i (Figs 3E, F and S3B). Taken together, these data suggest that STAT1 and STAT5B are required for autophagy induction by all three types of IFNs.

**Fig 3.**
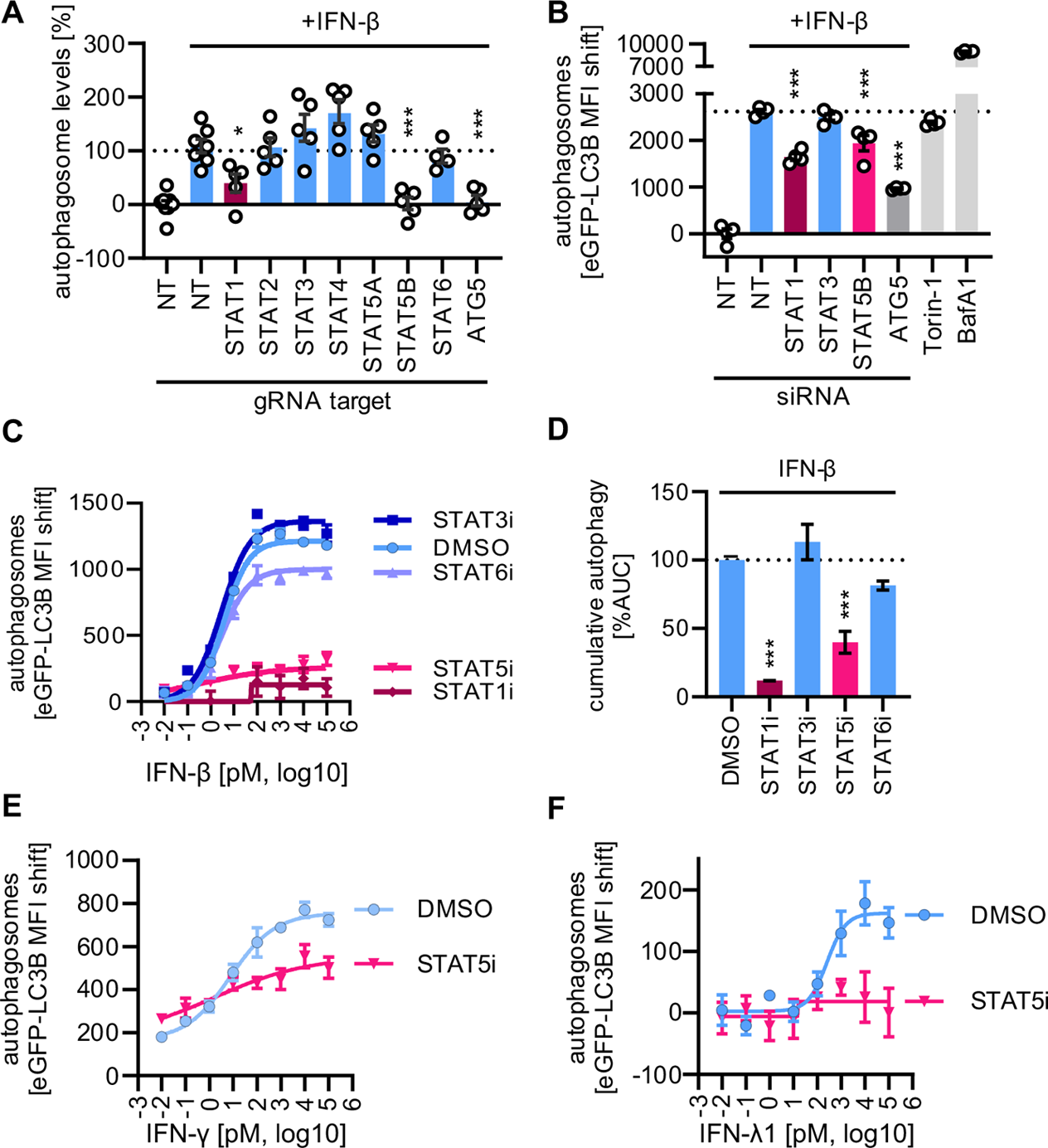
IFN induced autophagy is dependent on STAT1/5B. **A,** Quantification of autophagosome levels by flow cytometry in HEK293T autophagy reporter cells (HEK293T GL) transiently expressing Cas9 and at least 2 individual gRNAs against indicated STATs. Treated with 10 pM IFN-β for 24h. n = 4-6± SEM. **B,** Quantification of autophagosome levels by flow cytometry in HeLa autophagy reporter cells (HeLa GL) transfected with siRNAs targeting *STAT1*, *STAT3*, *STAT5B* or *ATG5* and treated with 1 nM IFN-β for 24h. BafA1 (625 µM) or Torin-1 (1 µM) for 4h used as controls. n = 4 ± SEM. **C**, Quantification of autophagosome levels by flow cytometry in HeLa autophagy reporter cells (HeLa GL) 24 h after treatment with increasing concentrations of IFN-β in presence or absence of 1 µM 5,15-DPP (STAT3 Inhibitor, STAT3i), 1 µM AS1517499 (STAT6 Inhibitor, STAT6i), 10 µM Fludarabine (STAT1 Inhibitor, STAT1i) or 100 µM STAT5 Inhibitor (STAT5i). n = 4 ± SEM. **D,** Area under the curve (AUC) analysis of the data in (C). **E**, Quantification of autophagosome levels by flow cytometry in HeLa autophagy reporter cells (HeLa GL) 24 h after treatment with increasing concentrations of IFN-γ and STAT5 Inhibitor (STAT5i, 100 µM). n = 4 ± SEM. **F**, Quantification of autophagosome levels by flow cytometry in HeLa autophagy reporter cells (HeLa GL) 24 h after treatment with increasing concentrations of IFN λ1 and STAT5 Inhibitor (STAT5i, 100 µM). n = 3 ± SEM. Brown-Forsythe and Welch ANOVA with Dunnett’s T3 multiple comparisons test. *, p<0.05, ** p<0.01, *** p<0.001.

### STAT1 and STAT5B form a complex that mediates autophagy induction

Our data indicated that depletion of STAT1 and STAT5B almost abolished IFN-induced autophagy. To examine whether both proteins work together, we performed overexpression studies. While individual expression of STAT1 and STAT5B only slightly induced autophagosomes, their combined overexpression significantly boosted autophagosome levels (Fig 4A), suggesting that the two proteins work synergistically. Expression of TRIM32 served as positive control [36,38,39]. To induce transcription of ISGs, STAT molecules typically form hetero- or homodimers [40,41]. Co-immunoprecipitation of overexpressed FLAG-STAT5B, FLAG-STAT1 and respective V5-tagged STAT1 or STAT5B revealed that both STAT1 and STAT5B bind to themselves as well as to one another with similar efficiency (Fig 4B). These interactions were confirmed at an endogenous level in primary NHLF cells, where STAT1 readily co-purified with STAT5B (Fig S4A). In line with this, proximity ligation assays in primary human dermal fibroblasts showed that upon treatment with IFN-β, STAT1 and STAT5B are in close spatial proximity (Figs 4C and S4B), suggesting that STAT1 and STAT5B associate after IFN treatment. To induce transcription, the STAT transcription factors translocate to the nucleus. Upon IFN-β stimulation, localization of endogenous STAT1 and STAT5B in primary NHLF cells significantly shifted to the nucleus (Figs 4D, E), where they co-localize as indicated by an increased Pearson correlation coefficient (Fig S4C). Taken together, these assays indicate that STAT1-STAT5B associate upon IFN stimulation, and traverse to the nucleus where they reside in close proximity.

**Fig 4.**
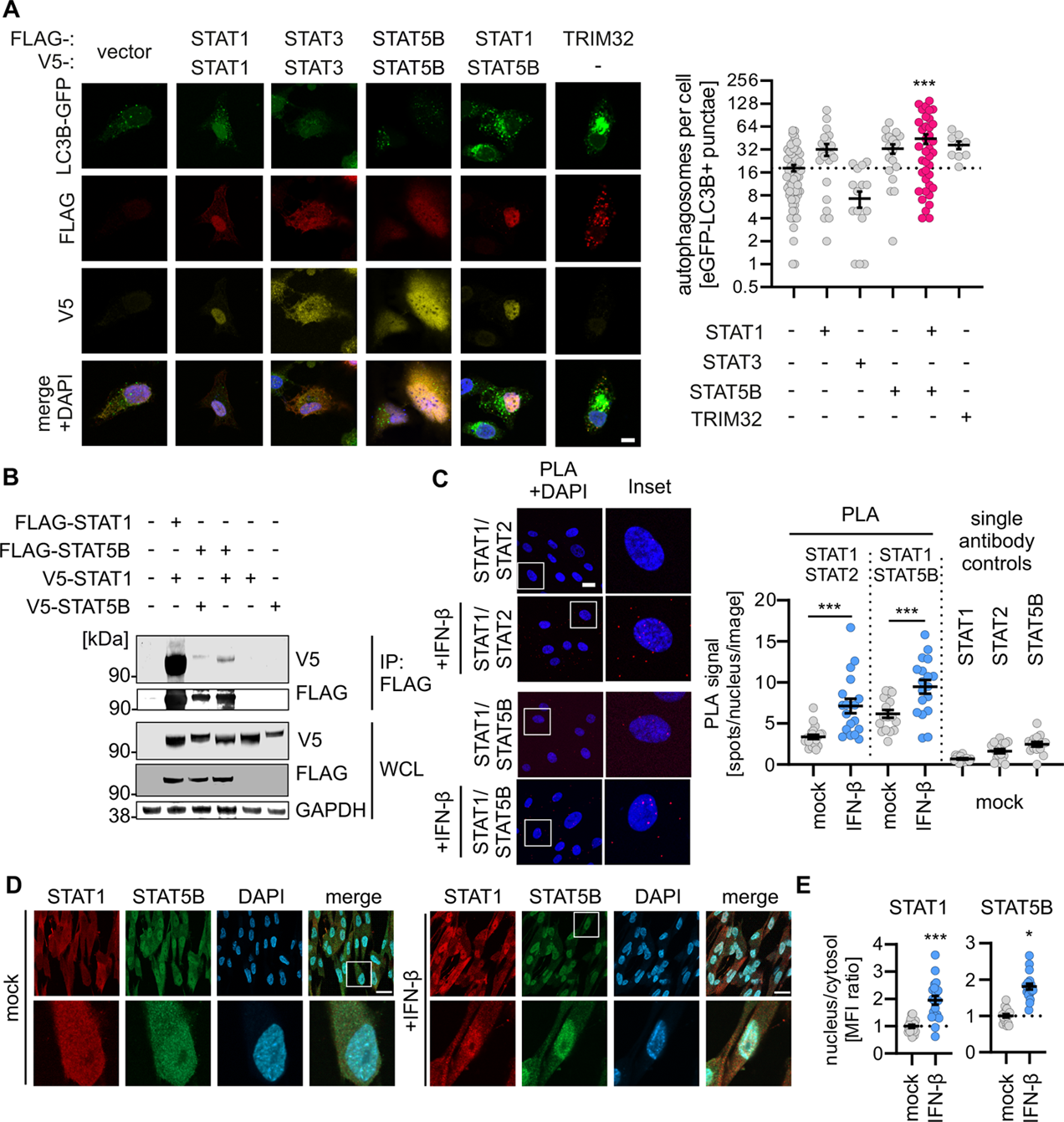
STAT1 and STAT5B interact to induce autophagy. **A,** Representative confocal immunofluorescence images of HeLa autophagy reporter cells (HeLa GL) transiently expressing FLAG-tagged (red) and V5-tagged (yellow) STAT1, STAT3 or STAT5 in combination or FLAG-tagged TRIM32 (scale bar, 10 μm) (left panel). Quantification of the number of autophagosomes (= eGFP-LC3B positive puncta) per cell in the images in the left panel. n= 9-63 cells ± SEM Brown-Forsythe and Welch ANOVA with Dunnett’s T3 multiple comparisons test. **B,** Co-immunoprecipitation of FLAG-tagged STAT1 and STAT5B from whole cell lysates of HEK293T cells transiently expressing FLAG-tagged and V5-tagged STAT1 and STAT5B or V5-tagged STAT1 or STAT5B alone. Immunoblots stained with anti-V5, anti-FLAG or anti-GAPDH. **C,** Exemplary images of Proximity Ligation Assay (PLA) of STAT1 and STAT2 or STAT5B (left panel) in HDF hTERT cells with or without treatment with 1 nM IFN-β for 1h. PLA signal (red). DAPI, nuclei (blue). Quantification of the number of PLA spots per nucleus per image of the left panel with single antibody controls (right panel). n = 15-21 tiles ± SEM. Student’s t-test with Welch’s correction. **D,** Representative confocal immunofluorescence images of NHLF cells treated with 1 nM of IFN-β for 1h and stained with α-STAT1 (green) and α-STAT5B (red), and DAPI (blue, nuclei) (scalebar 25 µm). **E,** Quantification of the ratio of nuclear to cytosolic intensity of STAT1 and STAT5B in the images in (D). n= 19-20 cells ± SEM. Student’s t-test with Welch’s correction. *, p<0.05, ** p<0.01, *** p<0.001

### SOCS1 is a STAT5B-dependent gene that drives autophagy induction

To examine which ISGs are induced in a STAT5 dependent fashion, we performed transcriptome analyses of IFN-β-treated HeLa cells in the presence and absence of the STAT5 inhibitor (Figs 5A and S5A, B; Supplementary Data File S1). Quality controls confirmed the integrity of the dataset (Fig S5A). Correlation analyses revealed that inhibitor treatment did not change induction of the majority of genes (Fig 5B), among them are classical ISGs such as OAS proteins, MX1 or RIG-I (Fig S5C). However, the correlation analyses as well as principal component (PCA) analyses also suggested that a set of genes is differentially regulated (Fig 5B) upon STAT5 inhibition. Among these are CXCL10, which is more upregulated upon STAT5 inhibition, and IFI27, RSAD2, IFIT1 and SOCS1, which are less induced in the presence of the STAT5 inhibitor (Fig 5B). To quantitatively delineate which genes are regulated by STAT5, we performed a Deming regression and calculated the deviation (PC2) of each gene to the perfect correlation (Supplementary Data File S2, Fig 5C). These analyses revealed that SOCS1 and SLFN5 are the most downregulated genes in the presence of the STAT5 inhibitor (Fig 5D, E). Analysis of normalized counts shows that inhibition of STAT5 reduces SOCS1 expression by ~15-fold, whereas the expression of SLFN5 is reduced ~3.5-fold. Dependency of SOCS1 on STAT5 was confirmed by qPCR analysis in human primary dermal fibroblasts (Fig 5F). Finally, to determine whether SOCS1 drives IFN-induced autophagy, we used siRNAs in HeLa autophagy reporter cells to abrogate RSAD2, SOCS1, as well as STAT1, STAT5B and ATG5 as positive controls, and afterwards stimulated with IFN-β. Knockdown efficiency of the individual genes was over 90% (Fig S5C). Depletion of SOCS1 dramatically reduced autophagy levels upon IFN-β induction (Fig 5F). *Vice versa*, overexpression of SOCS1, but not RSAD2 led to higher GFP-LC3B positive autophagosome levels in HeLa autophagy reporter cells (Fig 5G). In addition, expression of SOCS1 in HEK293T autophagy reporter cells increased their autophagosome content significantly, as analyzed by flow cytometry (Fig 5H). In both assays, overexpression of TRIM32 was used as a positive control. Altogether, these data showed that STAT5-dependent induction of SOCS1 is required for autophagy activation by IFN.

**Fig 5.**
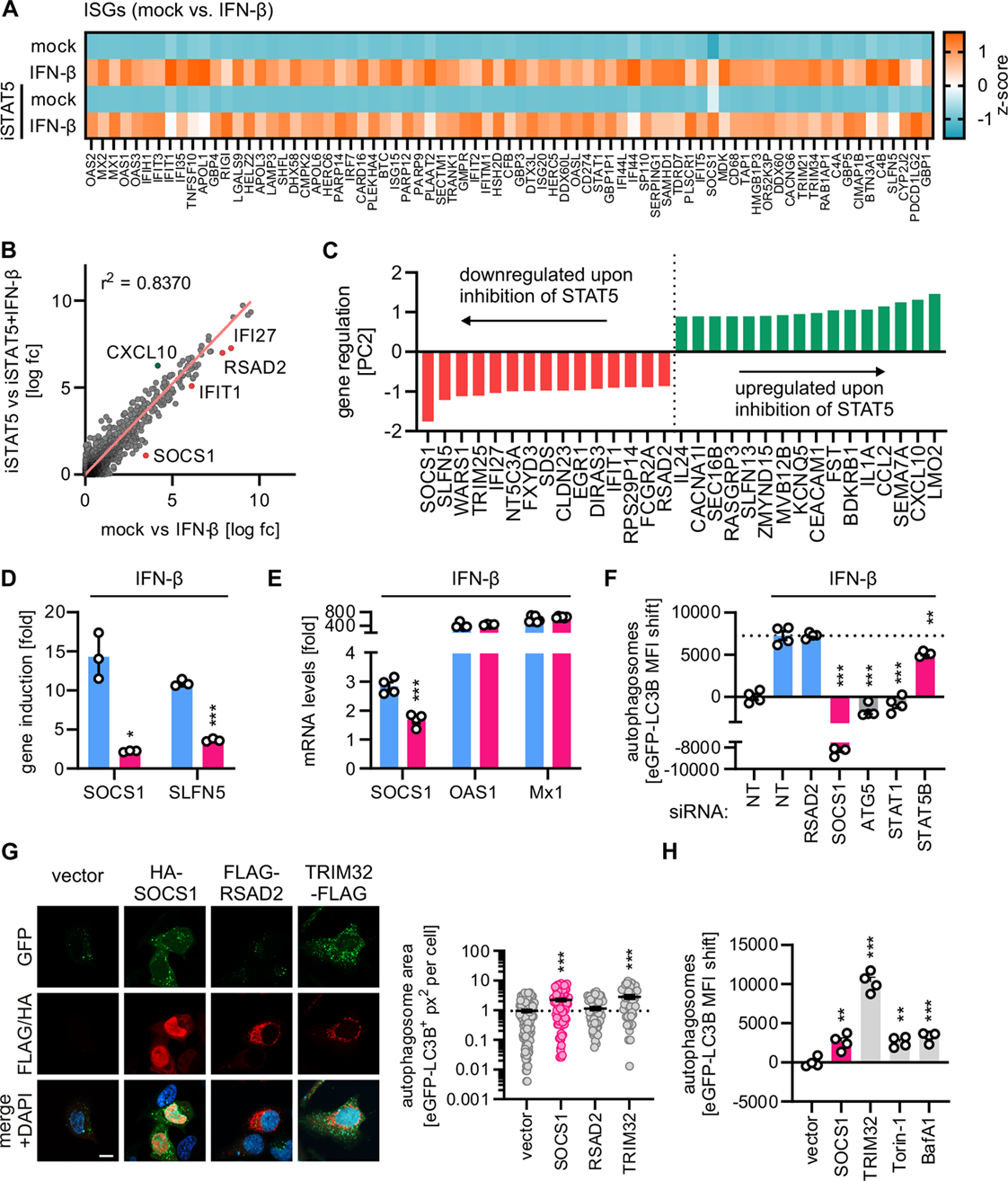
Identification of STAT5B-regulated genes responsible for autophagy induction. **A,** Heatmap of variance of the top 40 differentially expressed genes (DEGs) in HeLa GL cells treated with 1 nM IFN-β vs untreated control (mock vs. IFN-β) as assessed by next generation sequencing. Samples treated with 100 μM STAT5 Inhibitor are indicated. n = 3. **B,** Deming Correlation Analysis of log2-fold changes of DEGs (mock_vs_IFN-β and mock(iSTAT5)_vs_IFN-β (iSTAT5)) in HeLa GL in (A). Deming (Model II) Linear Regression. **C**, Quantification of Top20 up- and down-regulated genes by STAT5B (=PC2) of data in (B) by Principal Component Analysis. **D,** Fold changes of normalized counts of SOCS1 and SLFN5 derived from the data in (A and B). n=3± SEM. Student’s t-test with Welch’s correction. **E**, Quantification of gene expression by qRT-PCR of STAT5 regulated genes or ISGs in human dermal fibroblasts (HDF) cells treated with IFN-β (1 nM), STAT5i (250 μM) or both. Student’s t-test with Welch’s correction. **F,** Quantification of autophagosome levels by flow cytometry in HeLa autophagy reporter cells (HeLa GL) transfected with siRNAs for RSAD2, SOCS1, ATG5, STAT1 or STAT5B and treated with IFN-β (1 nM, 24h). n = 4 ± SEM. Brown-Forsythe and Welch ANOVA with Dunnett’s T3 multiple comparisons test. **(G),** Representative confocal immunofluorescence images of HeLa autophagy reporter cells (HeLa GL) transiently expressing FLAG- or HA-tagged (red) SOCS1, RSAD2 or TRIM32 (scale bar, 10 μm) (left panel). Quantification of the area of autophagosomes (= px^2^ of eGFP-LC3B positive puncta) per cell in the images in the left panel. n= 49-140 cells ± SEM. Brown-Forsythe and Welch ANOVA with Dunnett’s T3 multiple comparisons test. **H,** Quantification of autophagosome levels by flow cytometry in HEK293T autophagy reporter cells (HEL293T GL) expressing HA-SOCS1, TRIM32-FLAG or treated with Torin-1 (1µM, 4h) or Bafilomycin A1 (625nM, 4h) as positive controls. n= 4 ± SEM, Brown-Forsythe and Welch ANOVA with Dunnett’s T3 multiple comparisons test. *, p<0.05, ** p<0.01, *** p<0.001.

### Selective targeting of IFN-induced autophagy reduces growth of autophagy-dependent viruses

To determine the impact of the IFN-STAT1/STAT5B-SOCS1 axis on autophagy activation during viral infection, we infected HeLa autophagy reporter cells with Measles virus (MeV), Respiratory syncytial virus (RSV) and Influenza A virus (IAV). Inhibition of IFN-induced autophagy by STAT5i resulted in a drastic reduction of autophagosome levels as assessed by flow cytometry (Fig 6A). In line, autophagosome levels induced by infection in HeLa autophagy reporter cells were all significantly reduced upon inhibition of STAT5B, albeit to varying degrees (Figs 6B, C). Quantification of infection efficiency by immunofluorescence confirmed high infection rates (Fig S6). To dissect the impact of the IFN-autophagy axis on viral replication independently from ISG induction, we analyzed growth of MeV and RSV in the presence of IFN scavenging by B18R or STAT5i (Figs 6D, E). Expression of a GFP reporter from recombinant MeV and RSV was analyzed automatically every 4 hours in primary human lung fibroblasts (NHLF) to monitor virus growth. Cumulative gene expression analyses (area under the curve) showed that, as expected, scavenging IFN significantly promoted the growth of MeV (2.5-fold) and RSV (1.1-fold). In contrast, inhibition of the IFN-STAT5-SOCS1 axis restricted replication of MeV and RSV by 7.5- and 3.2-fold.

**Fig 6.**
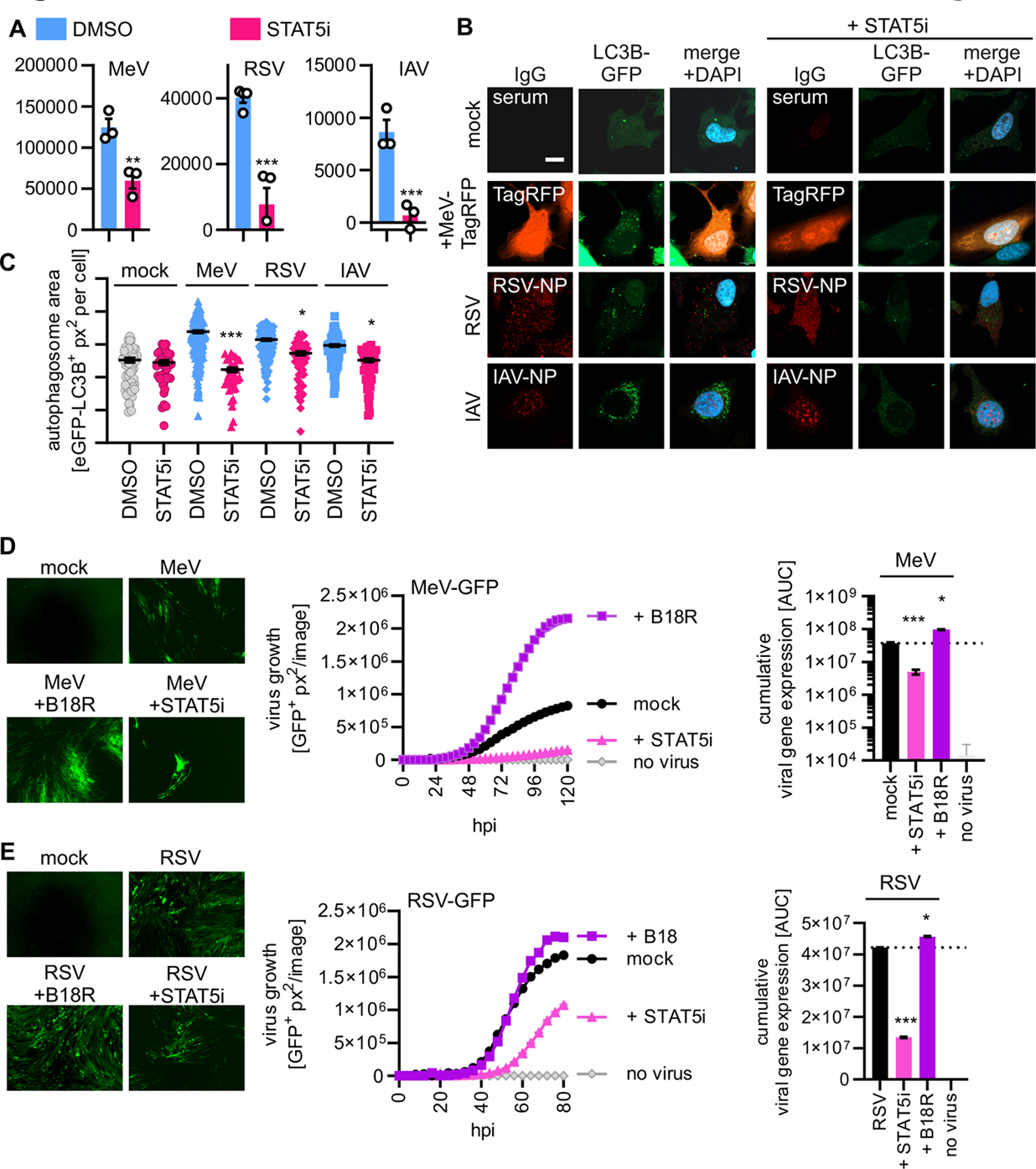
Virus-induced autophagy is dependent on IFN and STAT5B. **A,** Quantification of autophagosome levels by flow cytometry in HeLa autophagy reporter cells (HeLa GL) 24 h after infection with Measles Virus (MeV, MOI 1.25), Respiratory Syncytial Virus (RSV, MOI 1.25), or Influenza A Virus (IAV, MOI 1.25). Treated with 100 µM STAT5 Inhibitor (STAT5i) as indicated. n = 3 ± SEM. Student’s t-test with Welch’s correction. **B,** Representative confocal immunofluorescence images of HeLa autophagy reporter cells (HeLa GL, GFP-LC3B, green) 24 h after infection with Measles Virus (MeV, MOI 1), Respiratory Syncytial Virus (RSV, MOI 1), or Influenza A Virus (IAV, MOI 1) treated with STAT5i (100 µM, 24 h) as indicated. **C,** Quantification of the area of autophagosomes (= px^2^ of eGFP-LC3B positive puncta) per nucleus per tile of the images in (B). n= 25-50 tiles ± SEM. Brown-Forsythe and Welch ANOVA with Dunnett’s T3 multiple comparisons test. **D**, Exemplary microscopy images of NHLF cells infected with MeV-GFP (MOI 0.1) or left uninfected 3 days 16 h post infection in presence of 200 ng/ml B18R, 250 µM STAT5 Inhibitor (STAT5i) or without treatment (scale bar 100 μm, left panel). Quantification of viral growth by fluorescent reporter expression (= GFP-positive area (µm^2^)) over up to 6 days post infection of the images in the left panel (middle panel). n = 4-12 biological replicates ± SEM. Area-under the curve analysis of the growth curves in the middle panel (right panel). n = 4-12 biological replicates per group ± SEM. Brown-Forsythe and Welch ANOVA with Dunnett’s T3 multiple comparisons test. **E**, Exemplary microscopy images of NHLF cells infected with RSV-GFP (MOI 0.1) or left uninfected 3 days 4 h post infection in presence of 200 ng/ml B18R, 250 µM STAT5 Inhibitor (STAT5i) or without treatment (scale bar 100 μm, left panel). Quantification of viral growth by fluorescent reporter expression (= GFP-positive area (µm^2^)) over up to 80 h post infection of the images in the left panel (middle panel). n = 4-12 biological replicates ± SEM. Area-under the curve analysis of the growth curves in the middle panel (right panel). n = 4-12 biological replicates per group ± SEM. Brown-Forsythe and Welch ANOVA with Dunnett’s T3 multiple comparisons test. *, p<0.05, ** p<0.01, *** p<0.001.

Taken together, STAT5B-dependent signaling contributes significantly to autophagy induction by respiratory viruses and uncoupling IFN-induced autophagy from ISG responses by inhibiting STAT5 restricts replication of the autophagy-dependent viruses MeV and RSV.

## DISCUSSION

Concerted innate immune activation is required for effective pathogen defense [2]. Here, we show that upregulation of SOCS1 by STAT1/STAT5B-dependent signaling is interconnecting the IFN system and autophagy. Our mechanistic data shows that both STAT1 and STAT5B are required for effective autophagy induction, and loss or inhibition of either of them almost abrogates IFN-induced autophagy (Fig 3). Importantly, targeting of STAT5 abrogates IFN-induced autophagy, inhibiting replication of autophagy-dependent viruses such as MeV and RSV. Our work thus allows to dissect the coordinated activation of autophagy and the IFN system during viral infection and reveal targets for inhibition of viral replication by impairing autophagy, without compromising the defensive power of the IFN system.

While STAT1 is well-established as a crucial transcription factors mediating ISG induction, STAT5 play a major role in the development of many hematopoietic lineages, including B cells and T cells [42]. Of note, selective loss of STAT5B function by mutations in humans is associated with severe forms of pulmonary disease, eczema, combined immunodeficiency, autoimmune disease and bacterial and/or viral infections, consistent with our proposed role of STAT5B in innate immunity [43–45]. STAT5B is highly similar to its paralog STAT5A; however, it differs in 12 amino acids at the C-terminus, but the functional consequence of this difference is not understood yet. We reveal that IFN-mediated autophagy is dependent on STAT5B, but not STAT5A, suggesting distinct roles for these two highly similar proteins. The existence of pathogenic STAT5B mutations in humans also indicates that the functions of STAT5A and 5B are not redundant [46]. Moreover, STAT5B, but not STAT5A, was reported to have a key role in the development of leukemia [47] and has emerged as a novel target in pancreatic cancer [48]. Thus, our data, uncovers a so far unknown role of STAT5B in coordinating anti-viral innate immune responses, which is not redundant to the highly similar STAT5A.

The identity of the specific ISGs that require STAT5 as a transcription factor were previously unclear. It was suggested that STAT5 likely mediates induction of genes regulated by the (IFN) gamma activated sequence (GAS) promoter element [49–51] or that it may serve as a negative regulator of STAT1-mediated gene expression in cancers [52]. Our data now unveiled that STAT5 positively regulates transcription of a highly defined subset of ISGs, among them RSAD2 and SOCS1. Of note, induction of well-known anti-viral ISGs such as Mx1 or OAS1 is largely independent of STAT5B.

Autophagy regulation occurs at multiple levels including transcriptional and post-transcriptional regulation [53]. Our data shows that IFN induces autophagy, by activating STAT5-dependent transcription of SOCS1. SOCS proteins are well-known for acting as a negative feedback regulator of cytokines and being required for T cell homeostasis [54,55]. In line, genetic deletion of SOCS1 is lethal in mice due to systemic inflammation [56]. It was proposed that it may act as a direct inhibitor of JAKs [57] or exert their anti-inflammatory function via the recruitment of E3 ligases [58]. The contribution of SOCS1-induced autophagy to T cell homeostasis as well as the precise molecular mechanism of SOCS1 in autophagy, will be the subject of future studies. However, our data already uncovers a previously underappreciated axis of transcriptional regulation of autophagy.

Previous studies suggested that autophagy is induced by viral infections e.g. via activation of pattern recognition receptors like cGAS or by cellular stress responses [17,59,60]. Our data shows that during respiratory viral infection autophagy is boosted by IFN-STAT5B signaling 24 h post infection. This suggests that besides initial and rapid autophagy activation via PRRs or stress a second more sustained autophagy response to infection is induced.

In summary, we identified and characterized the IFN-STAT5B-SOCS1 axis as a key axis coordinating virus-induced autophagy, establishing a molecular link between the IFN system and autophagy. Selective targeting SOCS1 or STAT5B-dependent signaling may be a future approach to inhibit pro-viral autophagy while leaving other innate defenses intact.

## Materials and Methods

### Cell culture

The construction of autophagy reporter HEK293T and HeLa cells stably expressing GFP-LC3B (GL) was reported previously [36]. HEK293T (ATCC, #CRL321), HEK293T GL, Hela (ATCC, #CCL-2), HeLa GL, MDCK (ATCC, CCL-34), Vero E6 (ATCC, CRL-1586), HEp-2 (ATCC, # CCL-23), and A549 (ATCC, #CCL-185) cell lines and human dermal fibroblast (HDF) cells immortalized by stably expressing human telomerase reverse transcriptase (hTERT) (HDF hTERT) (provided by Patrick Hearing, Stony Brook University, New York, USA) as well as primary normal human lung fibroblast (NHLF, Lonza) were cultivated in Dulbecco’s Modified Eagle Medium (DMEM, gibco) supplemented with 10% (v/v) fetal bovine serum (FBS), 6.5 µg/ml (13 µM) gentamicin, and 2 mM L-glutamine. All cells were incubated at 37°C in a 5% CO_2_, 90% humidity atmosphere.

### Viruses

Influenza A Virus (A/PR/8/34 H1N1) was purchased from ATCC (VR-95). Measles virus (vac2), Measles virus expressing GFP (MeV-GFP, vac2) and Respiratory Syncytial Virus (A2) were a kind gift from Karl-Klaus Conzelmann. Respiratory Syncytial Virus expressing GFP (RSV-GFP, A2) was created by Michel N. Teng and provided by Kariem Ezzat Ahmed.

### Virus stock generation

Influenza A virus was grown as previously described [61,62]. In brief, MDCK cells were seeded in 75 cm^2^ cell culture flasks and incubated overnight. The next day, growth medium was removed and the cells were washed three times to remove any traces of FCS before infection with an MOI of 0.01. Infection and subsequent virus growth were carried out in 1 ml of cDMEM (DMEM supplemented with 6.5 µg/ml (13 µM) gentamicin, 2 mM L-glutamine, 0.2% Bovine Serum Albumin (BSA) and 25 mM HEPES) with L-(tosylamido-2-phenyl) ethyl chloromethyl ketone (TPCK) treated trypsin (1 µg/ml) added. The inoculum was removed after 1 h and the cells were washed once before the addition of 15 ml cDMEM. At 80% visible CPE, usually 2-3 days post-infection, medium and cells were harvested together by detaching the cells using a cells scraper, freezing the flask at −80°C and after thawing transfer the virus-containing supernatant to a 50 ml falcon. The supernatant was then sonicated for 10 min and cleared by centrifugation at 300 x g for 15 min at 4°C to pellet cell debris. Cleared supernatant was aliquoted and frozen at −80°C. Titer was determined by plaque assay as previously described [63]. In brief, MDCK cells were seeded in 12 well plates and infected with 100 µL of increasing dilutions, ranging from 10^1^ to 10^10^, with produced virus stocks in triplicates and 250 µL cDMEM. Inoculum was allowed to attach for 1 h, before removal and addition of 2 ml of overlay medium (cDMEM supplemented with 1% DEAE dextran, 5% NaHCO3, 1 µ/ml TPCK trypsin and 0.6% Avicel). 48-72 h after infection, the overlay was removed and cells were fixed using 4% PFA for 1 h at RT and washed with PBS. Plates were stained using 0.5% crystal violet solution in 30% EtOH for 10 min at RT. The staining solution was removed and the plates were washed three times. Plaques were counted after the plates were dried overnight.

Measles virus was grown on Vero E6 cells. Vero E6 cells were seeded into 75 cm^2^ cell culture flasks and incubated overnight. The next day, medium was removed and the cells were infected with an MOI of 0.01 in DMEM supplemented with 10% (v/v) fetal bovine serum (FBS), 6.5 µg/ml (13 µM) gentamicin, and 2 mM L-glutamine. 3-5 days post infection, at approximately 80% visible CPE, virus was harvested by removing the supernatant and adding 5 ml OptiMEM (Gibco) to the flask. The cells were frozen at −80°C for 20 min and scraped off the cell culture flask after thawing. The resulting virus-containing supernatant was sonicated for 10 min and cleared by centrifugation at 300 x g for 10 min at 4°C to pellet cell debris. Cleared supernatant was aliquoted and frozen at −80°C. For titration, Vero cells were seeded in 96 well plates and incubated overnight. The next day, cells were infected using 100 µL of increasing dilutions, ranging from 10^1^ to 10^10^, with produced virus stocks in triplicates. 72 h after infection, cells were fixed using 80% ice-cold acetone for 20 min at 4°C. After removal of acetone, plates were allowed to dry for 20 min before addition of FITC-tagged anti-Measles antibody, clone 83KKII, (MAB8906F, Merck Millipore) for 2 h at 37°C. Cells were washed twice with PBS, and FITC-positive foci were counted using a fluorescence microscope (Leica) or a Cytation 3 (BioTek).

Respiratory Syncytial Virus was grown on HEp-2 cells. Cells were seeded in 15 cm cell culture dishes and incubated overnight. The next day, medium was removed and the cells were infected with an MOI of 0.01 in 10 ml of DMEM without supplements. 4 h post-infection, the inoculum was removed and 20 ml of DMEM supplemented with 10% (v/v) fetal bovine serum (FBS), 6.5 µg/ml (13 µM) gentamicin, and 2 mM L-glutamine was added. After 3-5 days, at approximately the first visible syncytia, medium was changed to 10 ml DMEM supplemented with 10% (v/v) fetal bovine serum (FBS), 6.5 µg/ml (13 µM) gentamicin, and 2 mM L-glutamine per dish. 6-8 h after medium reduction virus was harvested by removing the supernatant and scraping the cells in the remaining medium. Supernatant was cleared at 1000 x g for 5 min at 4°C. The scraped cells were collected and vortexed for 5 min before debris was cleared at 1000 x g for 5 min at 4°C. Supernatant from the dished and cleared supernatant from the cells was pooled, aliquoted and frozen at −80°C. Infectious virus yields of RSV were determined by seeding HEp-2 cell in 96well plates and incubating overnight at 37°C, 5% CO2. The next day, cells were infected with 100 µl of increasing viral dilutions, ranging from 10^1^ to 10^9^, and incubated for 3-5 days until clearly visible CPE was detectable. At that time, cells were fixed using 4% PFA for 1 h at RT and washed with PBS. Plates were stained using 0.5% crystal violet solution in 30% EtOH for 10 min at RT. The staining solution was removed and the plates were washed three times. Infected wells were counted after the plates were dried overnight and TCID50/ml was calculated using the Reed-Muench interpolation method [64].

### Expression constructs and cloning

Constructs coding for human STAT1 (eGFP STAT1 WT, Addgene #12301, kindly gifted from Alan Perantoni), STAT3 (pEGFP-N1-STAT3, Addgene #111934, kindly gifted from Geert van den Bogaart) and STAT5B (MAC_N_STA5B, Addgene #167800, kindly gifted from Markku Varjosalo) were obtained from Addgene. pTwist-FLAG was generated by removing the ORF of TRIM3 from pTwist_3x_FLAG_optTRIM (purchased from Twist Biosciences) as described previously (Klute et al.). pTwist-V5 was generated by linearizing pTwist-FLAG by PCR using the pTwist-FLAG fwd and rev primers from Table 1, amplifying the V5-gBlock (IDT) and inserting it into the linearized pTwist ‘null’ vector using Gibson Assembly (New England Biolabs). The ORFs of STAT1, STAT3 and STAT5B were amplified using the respective primers from Table 1 and then subcloned into pTwist-FLAG and pTwist-V5 vectors using Gibson assembly (New England Biolabs) to generate pTwist-FLAG-STAT1, -STAT3, -STAT5B, pTwist-V5-STAT1, -STAT3, -STAT5B. The vectors were linearized with AfeI and NheI restriction enzymes (New England Biolabs). pIRES-FLAG-TRIM32 was previously described [19,36].

**Table 1:**
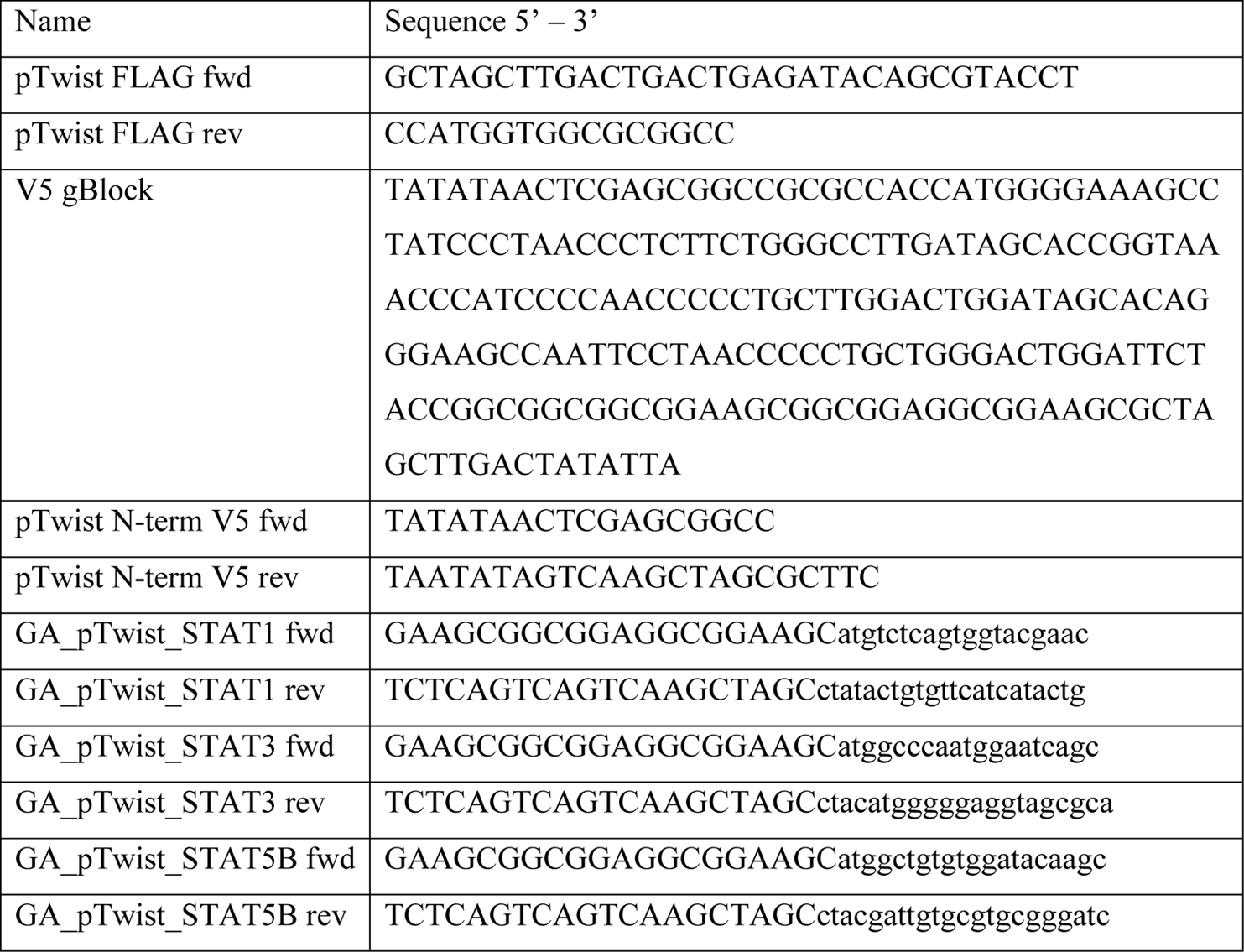
Primers used for cloning.

Constructs for CRISPR-Cas9-mediated knockout were generated by linearizing RP418 [65] with BsmbI (New England Biolabs), and single guideRNAs (Table 2) were inserted using Gibson Assembly (New England Biolabs). RP418 was a kind gift from Robert Jan Lebbink (University Medical Center Utrecht, Utrecht).

**Table 2:**
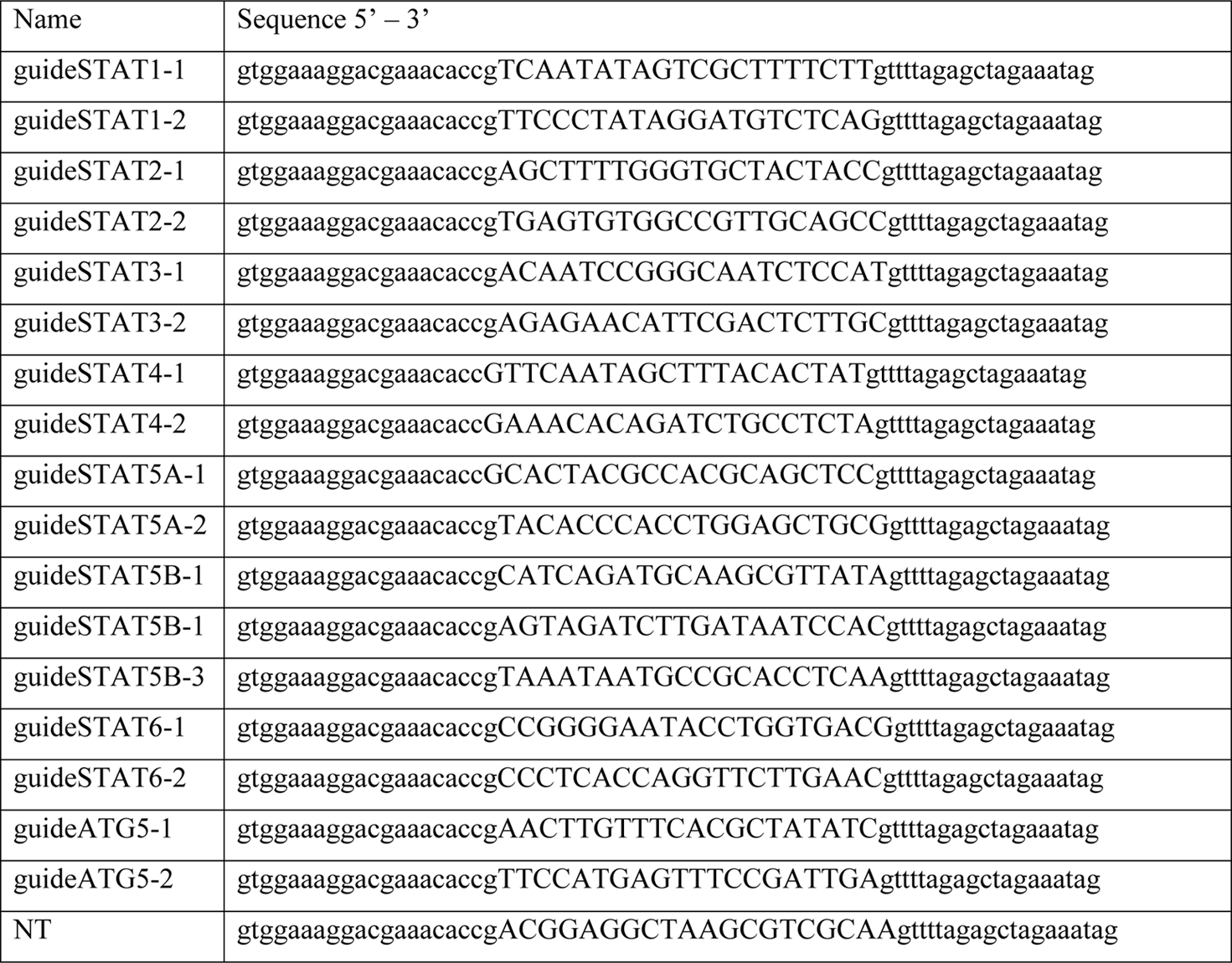
single guideRNAs (capital letters) with overhang (lowercase letters) used for cloning.

### Transfection of mammalian cells

For transfection, cells were seeded to reach a confluency of 60–80% the next day. Plasmid DNA was diluted in Opti-MEM (Gibco 31985070), and in parallel, PEI (Sigma-Aldrich; 408727) (2 µg PEI per 1 µg plasmid DNA) for transfection of HEK293T cells or TransIT LT1 (Mirus, MIR 2300) (3 µl transfection reagent per 1 µg plasmid DNA) for overexpression in HeLa GL cells was suspended in Opti-MEM. Both suspensions (plasmid in OptiMEM and transfection reagent in OptiMEM) were mixed in a ratio of 1:1. After incubation at RT for 20 min, the complete transfection mix was added to each well. The medium was replaced with fresh medium after 6–16 h to remove the transfection reagents.

### High-throughput Autophagy Reporter Assay

The autophagosome levels of HeLa or HEK 293T cells stably expressing LC3B-GFP were assessed as previously reported [36]. For interferon stimulation: The cells were grown in 96-well F-bottom plates and stimulated with increasing amounts of interferons (0.01 pM – 100 nM), Recombinant Human Interferon beta Protein (R&D System), Recombinant Human IFN-b (Peprotech), Human IFN Alpha A (Alpha 2a), carrier-free (pbl Assay Science), Recombinant Human IFN-gamma Protein (R&D Systems), Recombinant Human IL29/IFN-lambda 1 Protein (R&D Systems), Recombinant Human IFN-l1 (Peprotech), Recombinant Human IL28A/IFN-lambda 2 Protein (R&D Systems), Recombinant Human IL28B/IFN-lambda 3 Protein (R&D Systems), Recombinant Human IFN-lambda 4 Protein (R&D Systems), Recombinant Human IFN-epsilon Protein (R&D Systems)) for 4-48h. For samples that were treated with JAK Inhibitors (Ruxolitinib (R&D Systems), TC JL 37 (R&D Systems), PF06551600 malonate (R&D Systems) or CP 690550 citrate (Tofacitinib citrate) (R&D Systems)), STAT Inhibitors (Fludarabine (Cayman Chemical, Biomol), AS1517499 (Cayman Chemical, Biomol), 5,15-DPP (Cayman Chemical, Biomol), STAT5 Inhibitor – CAS 285986-31-4 (Calbiochem, Merck Millipore or Cayman Chemical, Biomol) or STAT5-in-1 (Target Molecules, Biomol)) or B18R (R&DSystems), the cells were pre-treated with the respective inhibitor for 1 h, and inhibitor was also added to medium at time of interferon treatment. For HEK293T GL cells overexpressing RP418, cells were seeded in 6-well plates and transfected with 2 ug of plasmid the next day. 6-16h post transfection a medium change was performed to remove the transfection complexes. 3 days after transfection, the cells were split to fresh 6-well plates and left to recover for further 3 days. Afterwards, the cells were seeded as before in 96-well F-bottom plates and incubated for 24 h before stimulation with 10 pM IFN-β for 24h. HEK293T GL cells for transient overexpression were transfected in reverse and seeded in 96-well F-bottom plates at a density of 40.000 cells per well. 16h post transfection the medium was replaced and the cells incubated for a total of 48h after transfection. For infection experiments using the high-throughput autophagy reporter assay, 40.000 HeLa GL cells were seeded in 96-well plates and incubated overnight. The next day, the cells were infected with a MOI of 1.25 PFU/ml for IAV, 1.25 FFU/ml for MeV, 1.25 TCID50/ml for RSV in 100 µl of DMEM, supplemented with 10% (v/v) fetal bovine serum (FBS), 6.5 µg/ml (13 µM) gentamicin, and 2 mM L-glutamine. For the uninfected controls, medium change was performed. For experiments using either B18R Interferon scavenger or STAT5 Inhibitor, the inhibitors were directly added to the infection medium and uninfected controls were treated with the same amount of inhibitor or left untreated. For knockdown using siRNA, the cells were seeded in 24-well plates and 24 h later transfected using RNAiMAX (Thermo Fisher). 24 - 48h later, the cells were treated with 100 pM IFN-β for further 24h. For samples that were treated with Bafilomycin A1 or Torin-1, 20 h after treatment, Bafilomycin A1 at a concentration of 625 μM or Torin-1 at a concentration of 1 nM was added to the medium and incubated for another 4 h. For flow cytometry preparation, the samples were then trypsinized and transferred to 96-well V-bottom plates. Cells transfected with siRNA were split and RNA from one half was isolated using the Quick-RNA Microprep Kit (Zymo research) according to the manufacturer’s instructions. Cells for autophagy assay were treated with 0.05% saponin in PBS and two subsequent washes with PBS were used to remove cytosolic LC3B-GFP. Fluorescence intensity of membrane-bound LC3B-GFP was measured using a BD FACSCanto II (BD Biosciences), a Beckman-Coulter CytoFLEX or a Cytek Northern Lights (Cytek Biosciences) with attached high-throughput sampler and baseline signal set to ≥1000 to allow for detection of shifts in autophagosome levels in both directions (more or less autophagosomes). Intact single cells were gates using SSC-A / FSC-A and FSC-A / FSC-H respectively. Raw data was analyzed using FlowJo 10. Median fluorescence intensity shifts of all samples were calculated by subtracting the LC3B-GFP-MFI of the appropriate control samples from the treated, transfected or infected samples.

### Whole-cell lysates

Whole-cell lysates (WCL) were prepared by harvesting cells in Phosphate-Buffered Saline (PBS, Gibco) and pelleting them by centrifugation (500 g, 4 °C, 5 min). The cell pellet was lysed using RIPA buffer (150 mM NaCl, 1% NP-40, 0.5% Deoxycholic acid, 0.1% sodium dodecyl sulfate, 50 mM Tris-HCl pH 8) by incubating on ice for 10 min with frequent vortexing at maximum speed for 30 s. Cell debris was removed by centrifugation (20,000 g, 4 °C, 20 min), and the protein concentration of the supernatants was quantified using a BCA assay (Pierce Rapid Gold BCA Protein Assay Kit, Thermo Fisher Scientific). The lysates were stored until analysis at −20 °C.

### BCA assay

BCA assay was done using the manufacturer’s instructions. In brief, 10 μl of lysates or albumin standard (Thermo Fisher Scientific) in a dilution series of 2 mg/ml – 0.125 mg/ml was added to clear F-bottom 96-well plates in duplicates. 200 μl of Working Reagent (50:1 mix of Reagent A and B) was added to the wells and incubated for 5 min at room temperature. Absorption at 480 nm was immediately measured on an Orion plate reader and concentrations were calculated using the standard curve.

### SDS-PAGE and Immunoblotting

SDS-PAGE and immunoblotting was performed as previously described[36]. In brief, whole cell lysates were mixed with 6x Protein Sample Loading Buffer (Tris-HCl pH 6.8, 75% Glycerol, 6% SDS, 0,3% w(v) Orange G, 15% β-mercaptoethanol) and heated to 96°C for 10 min, separated on NuPAGE 4-12% Bis-Tris Gels (Invitrogen) for 90-120 minutes at 90 V and transferred onto Immobilon-FL PVDF membranes (Merck Millipore) at a constant voltage of 30 V for 30 min. After the transfer, the membrane was dried overnight at room temperature and total protein staining was performed using the Revert 700 Total Protein Stain (LI-COR) according to the manufacturer’s instructions. After removal of the total protein stain, the membranes were blocked in 1% Casein in PBS. Proteins were stained with primary antibodies (mouse anti-SQSTM1/p62, abcam, #ab56416, 1:1000, mouse anti-FLAG M2, Sigma-Aldrich, #F1804, 1:3000, rabbit anti-V5, Cell Signaling, #13202, 1:1000, mouse anti-STAT1, Santa Cruz, #sc-464, 1:250, rabbit anti-STAT2, Cell Signaling, #72604, 1:250, rabbit anti-STAT5B, Genetex, #GTX08967, 1:250, rat anti-GAPDH, BioLegend, #607902, 1:3000) diluted in 0.1% Casein in PBS overnight at 4°C. Membranes were washed three times using PBS with 0.2% Tween20 and then stained with secondary IRDye-labelled secondary antibodies (LI-COR), diluted in 0.1% Casein in PBS for 1h at RT. Detection of proteins was done using either an Odyssey 9120 Imager (LI-COR) with the Image Studio software (Version 5.2, LI-COR) or an Odyssey M Imaging System (LI-COR) with the LI-COR Acquisition software (Version 2.2, LI-COR). Band intensities were quantified using Image Studio Lite (Version 5.0.21, LI-COR) or Empiria Studio (Version 3.2, LI-COR).

### Co-Immunoprecipitation

HEK293T cells were transfected with empty vectors (pTwist-FLAG and pTwist-V5), STAT1 (pTwist STAT1-FLAG and pTwist STAT1-V5 or pTwist STAT1-V5 alone), STAT5B (pTwist STAT5B-FLAG and pTwist STAT5B-V5 or pTwist STAT5B-V5 alone), or STAT1 and STAT5B (pTwist STAT1-FLAG or pTwist STAT1-V5 and pTwist STAT5B V5 or STAT5B FLAG, respectively). 24 h post transfection, WCLs were prepared and incubated with anti-FLAG M2 magnetic beads (Sigma-Aldrich) for 4 h at 4 °C on a rotating shaker. Subsequently, the beads were washed five times with RIPA buffer and incubated with 1x Protein Sample Loading Buffer supplemented with 15% β-mercaptoethanol. After heating to 95°C for 10 min the samples were separated by SDS-PAGE and then transferred to PVDF membranes for protein detection as described for immunoblotting.

### Immunofluorescence

NHLF or HeLa GL cells were seeded on coverslips (VWR) in 24-well plates and treated, transfected or infected as indicated. For treated cells, medium was removed and 500 µl medium containing either IFNs, Torin-1 or Bafilomycin A1 at the indicated concentrations was added and the cells were incubated for 24 h. For overexpression, cells were transfected using TransIT-LT1 and medium was replaced 6-16h post-transfection and cells were incubated for further 24 h. For infection, HeLa GL cells were infected with an MOI of 1 PFU/ml for IAV, 1 FFU/ml for MeV, or 1 TCID50/ml for RSV in 500 µl of DMEM, supplemented with 10% (v/v) fetal bovine serum (FBS), 6.5 µg/ml (13 µM) gentamicin, and 2 mM L-glutamine. For the uninfected controls, medium change was performed. For experiments using either B18R Interferon scavenger or STAT5 Inhibitor, the inhibitors were directly added to the infection medium and uninfected controls were treated with the same amount of inhibitor or left untreated. The cells were then incubated for 24 at 37°C with 5% CO_2_. Next, the samples were washed once with PBS and fixed in 4% paraformaldehyde solution (PFA) for 20 min at RT, then permeabilized and blocked with PBS containing 0.5 % Triton X-100 and 5 % FCS for 1 h at RT. Afterwards, the cells were washed with PBS and incubated overnight in a wet chamber at 4 °C with primary antibody (mouse anti-SQSTM1/p62, abcam, #ab56416, 1:200, mouse anti-FLAG M2, Sigma-Aldrich, #F1804, 1:400, rabbit anti-V5, 1:400, Cell Signaling, #13202, mouse anti-STAT1, 1:50 Santa Cruz, #sc-464, rabbit anti-STAT2, 1:50, Cell Signaling, #72604, rabbit anti-STAT5B, 1:50, Genetex, #GTX08967) diluted in PBS containing 1 % FCS. After washing with PBS containing 0.1 % Tween 20, the samples were incubated with the respective secondary antibodies (anti-mouse IgG (H+L) F(ab’)2 Fragment (Alexa Fluor 488 Conjugate), Cell Signaling, #4408, anti-mouse IgG (H+L) F(ab’)2 Fragment (Alexa Fluor 594 Conjugate), Cell Signaling, #8890, anti-mouse IgG (H+L) F(ab’)2 Fragment (Alexa Fluor 647 Conjugate), Cell Signaling, #4410, anti-rabbit IgG (H+L) F(ab’)2 Fragment (Alexa Fluor 488 Conjugate), Cell Signaling, #4412, anti-rabbit IgG (H+L) F(ab’)2 Fragment (Alexa Fluor 594 Conjugate), Cell Signaling, #8889, anti-rabbit IgG (H+L) F(ab’)2 Fragment (Alexa Fluor 647 Conjugate), Cell Signaling, #4414) and 500 ng/ml DAPI for 2 h at 4 °C in the dark. Next, the samples were washed three times with PBS containing 0.1 % Tween 20 and once with ultrapure water and the cover slips were mounted onto microscopy slides (VWR). Images were acquired using a Zeiss LSM 710 or Zeiss LSM 980 confocal laser scanning microscope with ZEN imaging software (Zeiss). Images were analyzed with ImageJ (Fiji). The number of SQSTM1/p62 positive particles and the particle size were analyzed using a custom ImageJ macro (Fiji). The nuclear and cytoplasmatic signal intensities were determined using the Analyze Tool in ImageJ (Fiji) on a 50×50 px field in the nuclear region as determined by DAPI staining or in the cytoplasmatic region as determined by exclusion of DAPI signal, but inclusion of either STAT1 or STAT5B positive region. Co-localization was determined using the Huygens Professional 19.10 software. Pearson coefficients were calculated with the “Huygens Colocalization Analyzer” applying automatic thresholds.

### Proximity Ligation Assay

Human derma fibroblasts hTERT cells were seeded on coverslips (VWR) in 24-well plates and treated with IFN-β for 60 min. Next, the samples were washed once with PBS and immediately fixed in 3.7% paraformaldehyde solution (PFA) for 10 min at RT, permeabilized with PBS containing 0.5 % Triton X-100 and for 7 min at RT and blocked with PBS containing 5% BSA for 1 h at RT. Afterwards, the cells were washed with PBS and incubated overnight in a wet chamber at 4 °C with primary antibody (mouse anti-STAT1, 1:50 Santa Cruz, #sc-464, rabbit anti-STAT2, 1:50, Cell Signaling, #72604, rabbit anti-STAT5B, 1:50, Genetex, #GTX08967) diluted in PBS containing 5 % BSA. Afterwards, the slides were washed and immediately stained using DuoLink In-situ Detection Reagents Far Red (Sigma) according to the manufacturer’s protocol. Images were acquired using a Zeiss LSM 980 confocal laser scanning microscope with ZEN imaging software (Zeiss). Images were analyzed with ImageJ (Fiji). The number of ligated spots and number of nuclei per tile was analyzed using a custom ImageJ macro (Fiji).

### Automated virus growth analyses

10.000 NHLF cells were seeded into 96-well plates. The next day, cells were infected with either MeV-GFP or RSV-GFP with a MOI of 0.1 in presence or absence of either 200 ng/ml B18R or 250 µM STAT5i or left uninfected in 180 µl DMEM, supplemented with 10% (v/v) fetal bovine serum (FBS), 6.5 µg/ml (13 µM) gentamicin, and 2 mM L-glutamine. The cells were then incubated for up to 6 days at 37°C with 5% CO2 inside an Incucyte SX5 G/O/NIR with 5 phase and green channel images per well taken every 4 hours. Viral growth was observed using the expression of the fluorescent reporter and total GFP positive area (µm^2^) was used as proxy for viral replication.

### RNASeq

HeLa GL cells were pre-treated with 100 µM STAT5 Inhibitor for 1 hour or left untreated and then stimulated with 1 nM IFN-β or left unstimulated. Total RNA was extracted 16 h post-stimulation using the Quick-RNA Miniprep Kit (Zymo research) according to the manufacturer’s instructions. Construction of stranded RNA libraries was performed using the sparQ RNA-Seq HMR Kit (QuantaBio) with sparQ PureMag Beads (QuantaBio) and sparQ UDI Adapters (QuantaBio) according to the manufacturer’s instructions. In brief, 1000 ng of isolated RNA was used as input for the fragmentation and depletion reaction using 4 µl of the proprietary Frag Prime RG depletion mix in 10 µl total reaction volume. The reaction was mixed by pipetting 5 times up and down and briefly centrifuging before incubation at 94°C for 8 min and then stepwise reduction of 5°C to 75-55°C every two minutes and then incubation at 37°C for 5 min and 25°C for 5 min. First strand synthesis was done using 4 µl of the proprietary 1st Strand Enzyme mix of the kit and adding it to the fragmentation and depletion reaction together with 6 µl nuclease-free water for 20 µl total reaction volume. 1st strand synthesis was done by incubating the reaction at 25°C for 10 min, followed by incubation at 42°C for 15 min and 70°C for 15 min. The reaction was then immediately placed on ice and the second strand synthesis was performed by adding 20 µl of the 2nd Strand Buffer and 10 µl of the 2nd Strand Enzyme Mix to the 1st Strand Synthesis Reaction to achieve a total reaction volume of 50 µl. The second strand synthesis reaction was then incubated at 16°C for 30 min, followed by an incubation at 62°C for 10 min. The finished reaction was then cleaned up using 90 µl of the sparQ PureMag beads per reaction. After cDNA purification, UDI adapters were ligated to the cDNA using the sparQ UDI Adapters. For this, 5 µl of 1:10 or 1:20 unique diluted UDI adapters together with 20 µl of Rapid Ligation Buffer and 10 µl of T4 Ligase were added to the cDNA product and incubated at 20°C for 15 min. Ligated product was purified using 70 µl of the sparQ PureMag beads. 23.5 µl of the ligated product was used for library amplification. 25 µl of HiFi Plus Master Mix, 1.5 µl of proprietary Primer Mix and the 23.5 µl purified, adapter-ligated DNA was mixed for a total reaction volume of 50 µl and incubated for 10 min at 37°C, followed by 45 sec at 98°C and 10-12 cycles of 20 sec at 98°C, 30 sec at 60°C and 30 sec at 72°C before an incubation for 1 min at 72°C. The amplified library was purified three times using 45 µl of the sparQ PureMag beads. Quality control and quantification of Input RNA and libraries was done using a TapeStation 4150 (Agilent) and RNA ScreenTape (Agilent) or D1000 ScreenTape (Agilent) respectively. Sample libraries were sequenced using a Illumina NextSeq. Analysis of sequencing results was performed using a custom pipeline consisting of FastQC [66] (version 0.11.9), BEDtools [67] (version 2.30.0) and RSeQC [68] (version 5.0.1) on R (version 4.2.2) for quality control, Trim Galore! (version 0.6.7) for adapter trimming, a custom tool for paired-end matching (Alexander Graf, LAFUGA), STAR [69] (version 2.7.10b) for mapping to the reference genome (GRCH38.p13) with default settings, featureCounts [70] (subread version 2.0.3) for gene expression counting and DESeq2 [71] (version 1.34.0) on R (version 4.1.3) for differential gene expression on a Galaxy platform (version 24.1, hosted by the Laboratory for Functional Genome Analysis (LAFUGA)). Further analysis was done in R (version 4.4.1) on RStudio (Posit, version 2024.09.1) with the following packages attached: EnhancedVolcano (version 1.22.0), BiocManager (version 1.30.25), cluster (version 2.1.6), missMDA (version 1.19), factoextra (version 1.0.7), FactoMineR (version 2.11), pheatmap (version 1.0.12), tidyverse (version 2.0.0) and biomaRt (version 2.60.1).

### siRNA-mediated knock down

HeLa GL cells were transfected with siRNA (Horizon Discovery SMARTPOOL for STAT1, STAT3, STAT5B, ATG5, SOCS1, RSAD2 and Non-targeting Pool #2) using RNAiMax Transfection Reagent (Invitrogen) according to the manufacturer’s instructions. In brief, cells were seeded in 24 wells to reach a confluency of 60-80% at time of transfection. 5 pmol siRNA was used per 24 well and diluted in OptiMEM. 1.5 µl of RNAiMAX transfection reagent was also diluted in OptiMEM, and both solutions were mixed at a ratio of 1:1 before vortexing and incubation for 5 min at RT. 24 h after transfection, the medium was changed and the cells incubated for further 24h before treatment with 1 nM IFN-β. Afterwards, the cells were either harvested for high-throughput Autophagy Reporter Assay or for reverse transcription quantitative real time–PCR (RT-qRT-PCR). For RT-qRT-PCR, total RNA was extracted 24 h post treatment using the Quick-RNA Microprep Kit (Zymo Research) according to the manufacturer’s instructions.

### STAT5 target Gene Expression

HDF cells were treated with 250 µM STAT5 Inhibitor for 1h or left untreated. Afterwards, cells were stimulated with 1 nM IFN-β or left unstimulated. Total RNA was extracted 24 h post stimulation using the Quick-RNA Microprep Kit (Zymo Research) for RT-qRT-PCR of STAT5 regulated ISGs.

### RNA purification

RNA was isolated using either the Quick-RNA Microprep Kit (Zymo Research) for samples <10^6 cells or Quick-RNA Miniprep Kit (Zymo Research) for samples for samples <10^7 cells. In brief, cells were lysed in 300-600 µl of RNA Lysis buffer. Equal amount of 100% RNase-free ethanol (Roth) was added and lysate was transferred to spin columns. Columns were washed with 400 µl Wash buffer and afterwards DNase digest was performed by incubating the spin columns with 5 U of DNase I for 15 min at RT. Afterwards, columns were washed using 400 µl Prep buffer and twice with 400 µl and 700 µl Wash buffer. RNA was eluted in 15 µl nuclease free water.

### Reverse transcription quantitative real time–PCR (RT-qRT-PCR)

RT-qRT-PCR was performed using the Luna Universal Probe One-Step RT-qPCR Kit (New England Biolabs) on a Quant Studio 7 Pro Real-Time PCR System (Applied Biosystems) according to the manufacturer’s instructions. In brief, 7.5 µl Luna Universal Probe One-Step Reaction Mix (2X), 0.75 µl of Luna WarmStart RT Enzyme Mix (20X), 0.6 µl of housekeeping TaqMan Gene Expression Assay (GAPDH), 0.6 µl of target TaqMan Gene Expression Assay, 1-3 µl of isolated RNA and nuclease free water up to 15 µl total reaction volume was mixed and added to 96-well optical reaction plates and run on a Quant Studio 7 Pro Real-Time PCR System (Applied Biosystems) with the following cycling conditions: 10 min at 55°C for reverse transcription, 1 min at 95°C for initial denaturation, and 40 cycles of 10 seconds at 95°C denaturation followed by 60 seconds at 60°C extension step with plate read. TaqMan primer/probes for each individual target gene were purchased as premixed TaqMan Gene Expression Assays (Thermo Fisher Scientific) and added to the reaction (See probes in table 4). Expression levels for each target gene were calculated by normalizing to GAPDH expression levels using the ΔΔCT method.

**Table 3:** Primers used for TaqMan qPCR.

| Type | Primers | Assay ID |
| --- | --- | --- |
| TaqMan | STAT1 | Hs01013996_m1 |
| TaqMan | STAT3 | Hs00374280_m1 |
| TaqMan | STAT5B | Hs00560026_m1 |
| TaqMan | ATG5 | Hs00355494_m1 |
| TaqMan | SOCS1 | Hs00705164_s |
| TaqMan | RSAD2 | Hs00369813_m |
| TaqMan | OAS1 | Hs00973635_m1 |

### Quantification and statistical analysis

Statistical analyses were performed using GraphPad PRISM 10 (version 10.2.3). P-values were determined using Brown-Forsythe and Welch ANOVA with Dunnett’s T3 multiple comparisons test for multiple comparisons or a two-tailed Student’s t test with Welch’s correction for comparison of two datasets. Unless otherwise stated, data are shown as the mean of at least two biological replicates ± SEM. Significant differences are indicated as: *, p < 0.05; **, p < 0.01; ***, p < 0.001. Not significant differences are not indicated. Specific statistical parameters are specified in the figure legends.

### Data and materials availability

All data are available in the main text or the supplementary materials. RNA sequencing data was submitted to NCBI Gene Expression Omnibus (GEO, Accession code: GSE302359).

## Acknowledgments

We thank Daniela Krnavek, Martha Mayer, Kerstin Regensburger, Nicola Schrott, Regina Burger and Magdalena Weiß for additional excellent technical assistance. This study was supported by the German ministry for education and research grant IMMUNOMOD-01KI2014 (K.S.) and the German Research Foundation (CRC1279, SP 1600/7-1, SP 1600/9-1, KM 5/3-1, SP 1600/13-1).

## Supplementary information captions

Fig S1. IFN-α, IFN-ε and IFN-λ3 induce autophagic flux.

Fig S2. Inhibition of IFN-α, IFN-ε and IFN-λ3 induced autophagy by JAK Inhibitors.

Fig S3. KD efficiency and reduction of IFN induced autophagy by STAT5i.

Fig S4. STAT1 and STAT5B interact to induce autophagy.

Fig S5. Next-generation sequencing quality control, analysis and KD efficiency.

Fig S6. Infection efficiencies.

Data file S1. RNAseq analysis

Data file S2. PC2 quantification

## References

1. Akira S, Uematsu S, Takeuchi O. Pathogen recognition and innate immunity. Cell. 2006;124: 783–801. doi:10.1016/j.cell.2006.02.015

2. Sparrer KMJ, Gack MU. Intracellular detection of viral nucleic acids. Curr Opin Microbiol. 2015;26: 1–9. doi:10.1016/j.mib.2015.03.001

3. Koepke L, Gack MU, Sparrer KM. The antiviral activities of TRIM proteins. Curr Opin Microbiol. 2021;59: 50–57. doi:10.1016/j.mib.2020.07.005

4. Hornung V, Ellegast J, Kim S, Brzózka K, Jung A, Kato H, et al. 5’-Triphosphate RNA is the ligand for RIG-I. Science. 2006;314: 994–997. doi:10.1126/science.1132505

5. Pichlmair A, Schulz O, Tan CP, Näslund TI, Liljeström P, Weber F, et al. RIG-I-mediated antiviral responses to single-stranded RNA bearing 5’-phosphates. Science. 2006;314: 997– 1001. doi:10.1126/science.1132998

6. Ablasser A, Goldeck M, Cavlar T, Deimling T, Witte G, Röhl I, et al. cGAS produces a 2’-5’-linked cyclic dinucleotide second messenger that activates STING. Nature. 2013;498: 380–384. doi:10.1038/nature12306

7. Randall RE, Goodbourn S. Interferons and viruses: an interplay between induction, signalling, antiviral responses and virus countermeasures. J Gen Virol. 2008;89: 1–47. doi:10.1099/vir.0.83391-0

8. Platanias LC. Mechanisms of type-I- and type-II-interferon-mediated signalling. Nat Rev Immunol. 2005;5: 375–386. doi:10.1038/nri1604

9. Yetter A, Uddin S, Krolewski JJ, Jiao H, Yi T, Platanias LC. Association of the Interferon-dependent Tyrosine Kinase Tyk-2 with the Hematopoietic Cell Phosphatase *. Journal of Biological Chemistry. 1995;270: 18179–18182. doi:10.1074/jbc.270.31.18179

10. Lazear HM, Schoggins JW, Diamond MS. Shared and Distinct Functions of Type I and Type III Interferons. Immunity. 2019;50: 907–923. doi:10.1016/j.immuni.2019.03.025

11. Hu X, Li J, Fu M, Zhao X, Wang W. The JAK/STAT signaling pathway: from bench to clinic. Sig Transduct Target Ther. 2021;6: 1–33. doi:10.1038/s41392-021-00791-1

12. Kessler DS, Veals SA, Fu XY, Levy DE. Interferon-alpha regulates nuclear translocation and DNA-binding affinity of ISGF3, a multimeric transcriptional activator. Genes Dev. 1990;4: 1753–1765. doi:10.1101/gad.4.10.1753

13. Schoggins JW. Interferon-Stimulated Genes: What Do They All Do? Annu Rev Virol. 2019;6: 567–584. doi:10.1146/annurev-virology-092818-015756

14. Kluge SF, Sauter D, Kirchhoff F. SnapShot: antiviral restriction factors. Cell. 2015;163: 774–774.e1. doi:10.1016/j.cell.2015.10.019

15. Levine B, Mizushima N, Virgin HW. Autophagy in immunity and inflammation. Nature. 2011;469: 323–335. doi:10.1038/nature09782

16. Choi Y, Bowman JW, Jung JU. Autophagy during viral infection - a double-edged sword. Nat Rev Microbiol. 2018;16: 341–354. doi:10.1038/s41579-018-0003-6

17. Klute S, Sparrer KMJ. Friends and Foes: The Ambivalent Role of Autophagy in HIV-1 Infection. Viruses. 2024;16: 500. doi:10.3390/v16040500

18. Kim J, Kundu M, Viollet B, Guan K-L. AMPK and mTOR regulate autophagy through direct phosphorylation of Ulk1. Nature cell biology. 2011;13: 132. doi:10.1038/ncb2152

19. Hoenigsperger H, Koepke L, Acharya D, Hunszinger V, Freisem D, Grenzner A, et al. CSNK2 suppresses autophagy by activating FLN-NHL-containing TRIM proteins. Autophagy. 2024;20: 994–1014. doi:10.1080/15548627.2023.2281128

20. He C, Klionsky DJ. Regulation mechanisms and signaling pathways of autophagy. Annu Rev Genet. 2009;43: 67–93. doi:10.1146/annurev-genet-102808-114910

21. Gatica D, Lahiri V, Klionsky DJ. Cargo recognition and degradation by selective autophagy. Nat Cell Biol. 2018;20: 233–242. doi:10.1038/s41556-018-0037-z

22. Richetta C, Grégoire IP, Verlhac P, Azocar O, Baguet J, Flacher M, et al. Sustained Autophagy Contributes to Measles Virus Infectivity. PLOS Pathogens. 2013;9: e1003599. doi:10.1371/journal.ppat.1003599

23. Sparrer KMJ, Pfaller CK, Conzelmann K-K. Measles virus C protein interferes with Beta interferon transcription in the nucleus. J Virol. 2012;86: 796–805. doi:10.1128/JVI.05899-11

24. Grégoire IP, Rabourdin-Combe C, Faure M. Autophagy and RNA virus interactomes reveal IRGM as a common target. Autophagy. 2012;8: 36–1137. doi:10.4161/auto.20339

25. Chen L, Zhang J, Xu W, Chen J, Tang Y, Xiong S, et al. Cholesterol-rich lysosomes induced by respiratory syncytial virus promote viral replication by blocking autophagy flux. Nat Commun. 2024;15: 6311. doi:10.1038/s41467-024-50711-4

26. Gui X, Yang H, Li T, Tan X, Shi P, Li M, et al. Autophagy induction via STING trafficking is a primordial function of the cGAS pathway. Nature. 2019;567: 262–266. doi:10.1038/s41586-019-1006-9

27. Lee N-R, Ban J, Lee N-J, Yi C-M, Choi J-Y, Kim H, et al. Activation of RIG-I-Mediated Antiviral Signaling Triggers Autophagy Through the MAVS-TRAF6-Beclin-1 Signaling Axis. Front Immunol. 2018;9: 2096. doi:10.3389/fimmu.2018.02096

28. Delgado MA, Elmaoued RA, Davis AS, Kyei G, Deretic V. Toll-like receptors control autophagy. EMBO J. 2008;27: 1110–1121. doi:10.1038/emboj.2008.31

29. Yum S, Li M, Fang Y, Chen ZJ. TBK1 recruitment to STING activates both IRF3 and NF-κB that mediate immune defense against tumors and viral infections. Proceedings of the National Academy of Sciences. 2021;118: e2100225118. doi:10.1073/pnas.2100225118

30. Sparrer KMJ, Gableske S, Zurenski MA, Parker ZM, Full F, Baumgart GJ, et al. TRIM23 mediates virus-induced autophagy via activation of TBK1. Nat Microbiol. 2017;2: 1543–1557. doi:10.1038/s41564-017-0017-2

31. Du Y, Duan T, Feng Y, Liu Q, Lin M, Cui J, et al. LRRC25 inhibits type I IFN signaling by targeting ISG15-associated RIG-I for autophagic degradation. The EMBO Journal. 2018;37: 351–366. doi:10.15252/embj.201796781

32. Prabakaran T, Bodda C, Krapp C, Zhang B-C, Christensen MH, Sun C, et al. Attenuation of cGAS-STING signaling is mediated by a p62/SQSTM1-dependent autophagy pathway activated by TBK1. EMBO J. 2018;37: e97858. doi:10.15252/embj.201797858

33. Li P, Du Q, Cao Z, Guo Z, Evankovich J, Yan W, et al. Interferon-gamma induces autophagy with growth inhibition and cell death in human hepatocellular carcinoma (HCC) cells through interferon-regulatory factor-1 (IRF-1). Cancer Lett. 2012;314: 213–222. doi:10.1016/j.canlet.2011.09.031

34. Tian Y, Wang M-L, Zhao J. Crosstalk between Autophagy and Type I Interferon Responses in Innate Antiviral Immunity. Viruses. 2019;11: 132. doi:10.3390/v11020132

35. Symons JA, Alcamí A, Smith GL. Vaccinia virus encodes a soluble type I interferon receptor of novel structure and broad species specificity. Cell. 1995;81: 551–560. doi:10.1016/0092-8674(95)90076-4

36. Koepke L, Winter B, Grenzner A, Regensburger K, Engelhart S, van der Merwe JA, et al. An improved method for high-throughput quantification of autophagy in mammalian cells. Sci Rep. 2020;10: 12241. doi:10.1038/s41598-020-68607-w

37. Klionsky DJ, Abdel-Aziz AK, Abdelfatah S, Abdellatif M, Abdoli A, Abel S, et al. Guidelines for the use and interpretation of assays for monitoring autophagy (4th edition)1. Autophagy. 2021;17: 1–382. doi:10.1080/15548627.2020.1797280

38. You L, Wang Z, Li H, Shou J, Jing Z, Xie J, et al. The role of STAT3 in autophagy. Autophagy. 2015;11: 729–739. doi:10.1080/15548627.2015.1017192

39. Klute S, Nchioua R, Cordsmeier A, Vishwakarma J, Koepke L, Alshammary H, et al. SARS-CoV-2 Omicron Envelope T9I adaptation confers resistance to autophagy. bioRxiv; 2024. p. 2024.04.23.590789. doi:10.1101/2024.04.23.590789

40. Schindler U, Wu P, Rothe M, Brasseur M, McKnight SL. Components of a stat recognition code: Evidence for two layers of molecular selectivity. Immunity. 1995;2: 689–697. doi:10.1016/1074-7613(95)90013-6

41. Chen X, Vinkemeier U, Zhao Y, Jeruzalmi D, Darnell JE, Kuriyan J. Crystal Structure of a Tyrosine Phosphorylated STAT-1 Dimer Bound to DNA. Cell. 1998;93: 827–839. doi:10.1016/S0092-8674(00)81443-9

42. Heltemes-Harris LM, Willette MJL, Vang KB, Farrar MA. The role of STAT5 in the development, function, and transformation of B and T lymphocytes. Ann N Y Acad Sci. 2011;1217: 18–31. doi:10.1111/j.1749-6632.2010.05907.x

43. Nadeau K, Hwa V, Rosenfeld RG. STAT5b deficiency: an unsuspected cause of growth failure, immunodeficiency, and severe pulmonary disease. J Pediatr. 2011;158: 701–708. doi:10.1016/j.jpeds.2010.12.042

44. Cohen AC, Nadeau KC, Tu W, Hwa V, Dionis K, Bezrodnik L, et al. Cutting edge: Decreased accumulation and regulatory function of CD4+ CD25(high) T cells in human STAT5b deficiency. J Immunol. 2006;177: 2770–2774. doi:10.4049/jimmunol.177.5.2770

45. Vargas-Hernández A, Forbes LR. The Impact of Immunodeficiency on NK Cell Maturation and Function. Curr Allergy Asthma Rep. 2019;19: 2. doi:10.1007/s11882-019-0836-8

46. Klammt J, Neumann D, Gevers EF, Andrew SF, Schwartz ID, Rockstroh D, et al. Dominant-negative STAT5B mutations cause growth hormone insensitivity with short stature and mild immune dysregulation. Nat Commun. 2018;9: 2105. doi:10.1038/s41467-018-04521-0

47. Kollmann S, Grundschober E, Maurer B, Warsch W, Grausenburger R, Edlinger L, et al. Twins with different personalities: STAT5B—but not STAT5A—has a key role in BCR/ABL-induced leukemia. Leukemia. 2019;33: 1583–1597. doi:10.1038/s41375-018-0369-5

48. Moser C, Ruemmele P, Gehmert S, Schenk H, Kreutz MP, Mycielska ME, et al. STAT5b as Molecular Target in Pancreatic Cancer—Inhibition of Tumor Growth, Angiogenesis, and Metastases. Neoplasia. 2012;14: 915–925.

49. Wang W, Xu L, Su J, Peppelenbosch MP, Pan Q. Transcriptional Regulation of Antiviral Interferon-Stimulated Genes. Trends Microbiol. 2017;25: 573–584. doi:10.1016/j.tim.2017.01.001

50. Zhu M, John S, Berg M, Leonard WJ. Functional association of Nmi with Stat5 and Stat1 in IL-2- and IFNgamma-mediated signaling. Cell. 1999;96: 121–130. doi:10.1016/s0092-8674(00)80965-4

51. Platanias LC. Mechanisms of type-I- and type-II-interferon-mediated signalling. Nat Rev Immunol. 2005;5: 375–386. doi:10.1038/nri1604

52. Wellbrock C, Weisser C, Hassel JC, Fischer P, Becker J, Vetter CS, et al. STAT5 Contributes to Interferon Resistance of Melanoma Cells. Current Biology. 2005;15: 1629–1639. doi:10.1016/j.cub.2005.08.036

53. Lei Y, Klionsky DJ. Transcriptional regulation of autophagy and its implications in human disease. Cell Death Differ. 2023;30: 1416–1429. doi:10.1038/s41418-023-01162-9

54. Hilton DJ, Richardson RT, Alexander WS, Viney EM, Willson TA, Sprigg NS, et al. Twenty proteins containing a C-terminal SOCS box form five structural classes. Proc Natl Acad Sci U S A. 1998;95: 114–119. doi:10.1073/pnas.95.1.114

55. Ilangumaran S, Bobbala D, Ramanathan S. SOCS1: Regulator of T Cells in Autoimmunity and Cancer. Curr Top Microbiol Immunol. 2017;410: 159–189. doi:10.1007/82_2017_63

56. Starr R, Metcalf D, Elefanty AG, Brysha M, Willson TA, Nicola NA, et al. Liver degeneration and lymphoid deficiencies in mice lacking suppressor of cytokine signaling-1. Proc Natl Acad Sci U S A. 1998;95: 14395–14399. doi:10.1073/pnas.95.24.14395

57. Liau NPD, Laktyushin A, Lucet IS, Murphy JM, Yao S, Whitlock E, et al. The molecular basis of JAK/STAT inhibition by SOCS1. Nat Commun. 2018;9: 1558. doi:10.1038/s41467-018-04013-1

58. Babon JJ, Sabo JK, Zhang J-G, Nicola NA, Norton RS. The SOCS box encodes a hierarchy of affinities for Cullin5: implications for ubiquitin ligase formation and cytokine signalling suppression. J Mol Biol. 2009;387: 162–174. doi:10.1016/j.jmb.2009.01.024

59. Kroemer G, Mariño G, Levine B. Autophagy and the integrated stress response. Mol Cell. 2010;40: 280–293. doi:10.1016/j.molcel.2010.09.023

60. Gui X, Yang H, Li T, Tan X, Shi P, Li M, et al. Autophagy induction via STING trafficking is a primordial function of the cGAS pathway. Nature. 2019;567: 262–266. doi:10.1038/s41586-019-1006-9

61. Xue J, Chambers BS, Hensley SE, López CB. Propagation and Characterization of Influenza Virus Stocks That Lack High Levels of Defective Viral Genomes and Hemagglutinin Mutations. Front Microbiol. 2016;7: 326. doi:10.3389/fmicb.2016.00326

62. Eisfeld AJ, Neumann G, Kawaoka Y. Influenza A Virus Isolation, Culture and Identification. Nat Protoc. 2014;9: 2663–2681. doi:10.1038/nprot.2014.180

63. Matrosovich M, Matrosovich T, Garten W, Klenk H-D. New low-viscosity overlay medium for viral plaque assays. Virol J. 2006;3: 63. doi:10.1186/1743-422X-3-63

64. Reed LJ, Muench H. A Simple Method of Estimating Fifty Per Cent Endpoints12. American Journal of Epidemiology. 1938;27: 493–497. doi:10.1093/oxfordjournals.aje.a118408

65. van Diemen FR, Kruse EM, Hooykaas MJG, Bruggeling CE, Schürch AC, van Ham PM, et al. CRISPR/Cas9-Mediated Genome Editing of Herpesviruses Limits Productive and Latent Infections. PLoS Pathog. 2016;12: e1005701. doi:10.1371/journal.ppat.1005701

66. FastQC: A quality control tool for high throughput sequence data – ScienceOpen. [cited 9 May 2025]. Available: https://www.scienceopen.com/document?vid=de674375-ab83-4595-afa9-4c8aa9e4e736

67. Quinlan AR, Hall IM. BEDTools: a flexible suite of utilities for comparing genomic features. Bioinformatics. 2010;26: 841–842. doi:10.1093/bioinformatics/btq033

68. Wang L, Wang S, Li W. RSeQC: quality control of RNA-seq experiments. Bioinformatics. 2012;28: 2184–2185. doi:10.1093/bioinformatics/bts356

69. Dobin A, Davis CA, Schlesinger F, Drenkow J, Zaleski C, Jha S, et al. STAR: ultrafast universal RNA-seq aligner. Bioinformatics. 2013;29: 15–21. doi:10.1093/bioinformatics/bts635

70. Liao Y, Smyth GK, Shi W. featureCounts: an efficient general purpose program for assigning sequence reads to genomic features. Bioinformatics. 2014;30: 923–930. doi:10.1093/bioinformatics/btt656

71. Love MI, Huber W, Anders S. Moderated estimation of fold change and dispersion for RNA-seq data with DESeq2. Genome Biology. 2014;15: 550. doi:10.1186/s13059-014-0550-8

