## Supplementary Figure S1-S6 for "Respiratory viruses activate autophagy via the IFN-STAT1/STAT5B-SOCS1 axis"

#### Supplementary Figures S1-6

### Respiratory viruses activate autophagy via the IFN-STAT1/STAT5B-SOCS1 axis

Victoria Hunszinger[1], Helen Dürr[1], Zoé Engels[1], Helene Hönigsperger[1], Lennart Koepke[1], Susanne Klute[1], Jana-Romana Fischer[2], Birgit Ott[1], Alexander Graf[3], Stefan Krebs[3], Helmut Blum[3], Maximilian Hirschenberger[1], Frank Kirchhoff[1] and Konstantin MJ Sparrer[1,2*]

[1]Institute of Molecular Virology, Ulm University Medical Center, Ulm, Germany
[2]German Center for Neurodegenerative Diseases (DZNE), Ulm, Germany
[3]Laboratory for Functional Genome Analysis Gene Center, LMU Munich, Munich, Germany.

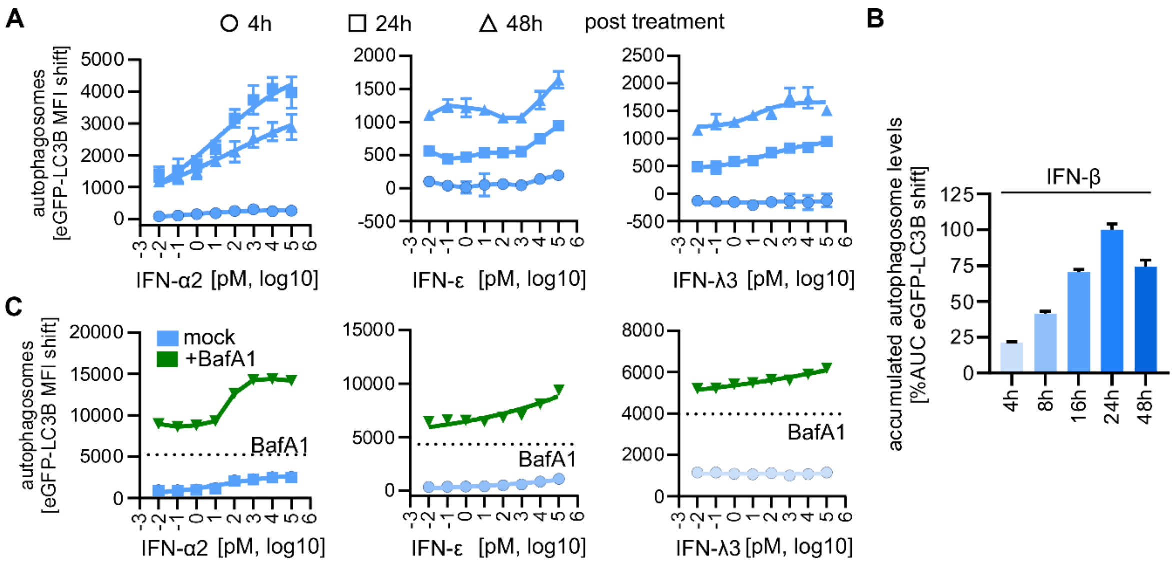


**Fig S1. IFNs induce autophagic flux. A,** Quantification of autophagosome levels by flow cytometry in HeLa autophagy reporter cells (HeLa GL) 4, 24 and 48 h after treatment with increasing concentrations (0.01 pM – 100 nM) of indicated IFNs. n = 4 ± SD. **B,** Area under the curve analysis of accumulated autophagy levels over time normalized by highest induction (AUC) of data in Fig 1a and additionally 8 and 16 h after treatment with IFN-β. n = 4-8 ± SEM. **C,** Quantification of autophagosome levels by flow cytometry in HeLa autophagy reporter cells (HeLa GL) 24 after treatment with increasing concentrations (0.01 pM – 100 nM) of indicated IFNs in presence of 625 nM of the autophagy flux inhibitor Bafilomycin A1. Dotted line, Bafilomycin A1 only treatment. n = 4 ± SD.

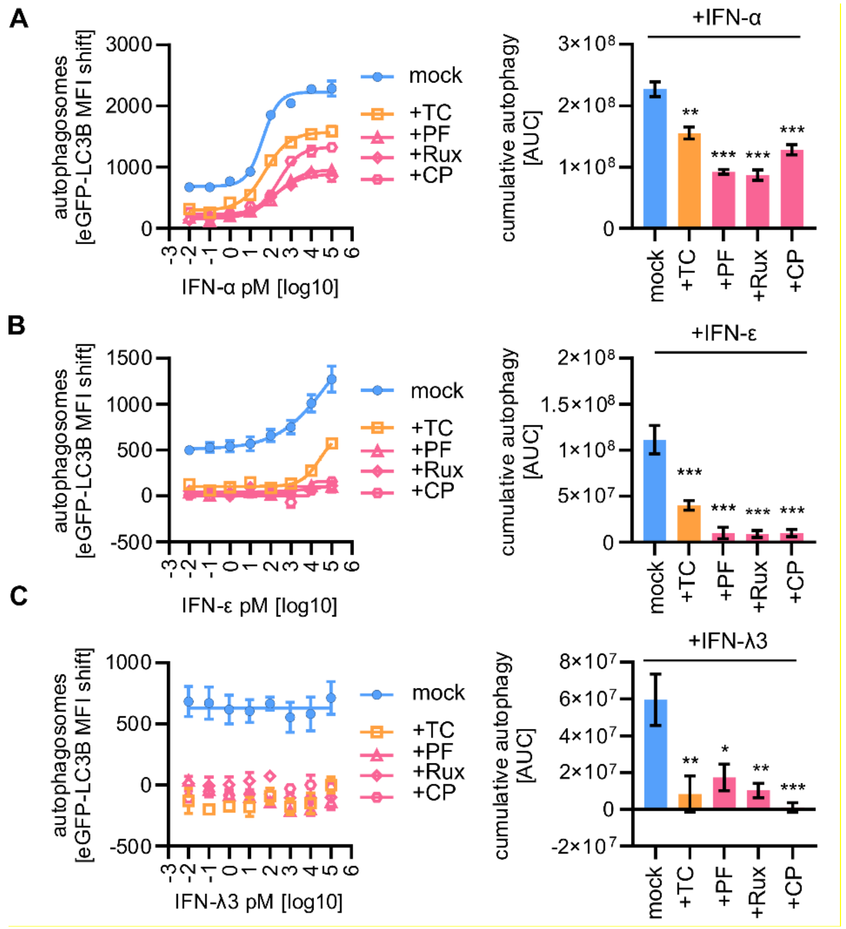


**Fig S2. Inhibition of IFN induced autophagy by JAK Inhibitors. A-C,** Quantification of autophagosome levels by flow cytometry in HeLa autophagy reporter cells (HeLa GL) 24 h after treatment with increasing concentrations (0.01 pM – 100 nM) of IFN-α (A), IFN-ε (B) or IFN-λ3 (C). Treated as indicated with TC JL 37 (TC, 100 nM), PF06551600 malonate (PF, Ritlecitinib, 100 µM), Ruxolitinib (Rux, 100 nM), or CP 690550 citrate (CP, Tofacitinib citrate, 100 nM), n = 4 ± SEM (left panels). Area under the curve (AUC) analysis of the data in (a-c**)**, (right panels). Ordinary one-way ANOVA with Dunnett's multiple comparisons test. * $p<0.05$, ** $p<0.01$, *** $p<0.001$.

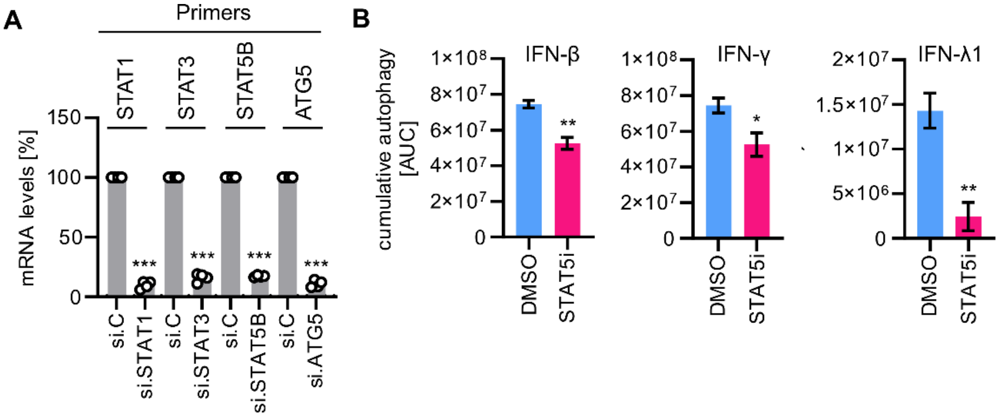


**Fig S3. KD efficiency and reduction of IFN induced autophagy by STAT5i. A,** RT-qRT-PCR analysis of siRNA-mediated knockdown of mRNA expression of data in Fig 3B. n = 4 ± SEM Student's t-test with Welch's correction. *** p<0.001. **B,** Area under the curve (AUC) analysis of the data in Figs 3E and 3F. n = 4 ± SEM Student's t-test with Welch's correction. ** p<0.01.

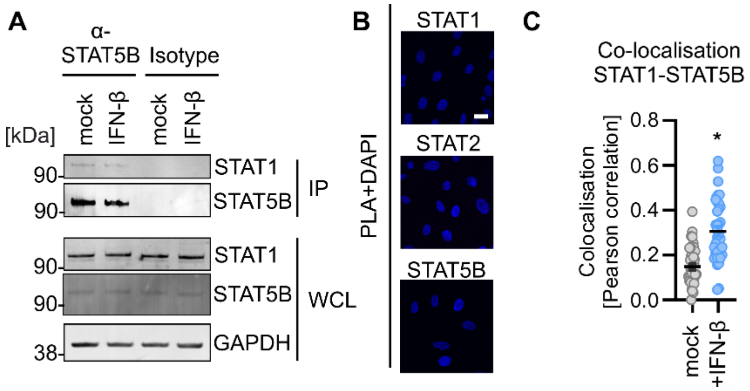


**Fig S4. STAT1 and STAT5B interact to induce autophagy. A,** Exemplary Immunoblot of the co-immunoprecipitation of endogenous STAT1 and STAT5B in NHLF cells treated with 1 nM IFN-β for 1h or left untreated. **B,** Exemplary confocal immunofluorescence images of single antibody controls for PLA in HDF hTERT cells of data in Fig 4c. PLA signal red. DAPI, nuclei (blue). (scale bar = 10 µm). **C,** Pearson Correlation of STAT1 and STAT5B signal of immunofluorescence data in Fig 4D. n = 37-38 ± SEM Student's t-test with Welch's correction. *** p<0.0001.

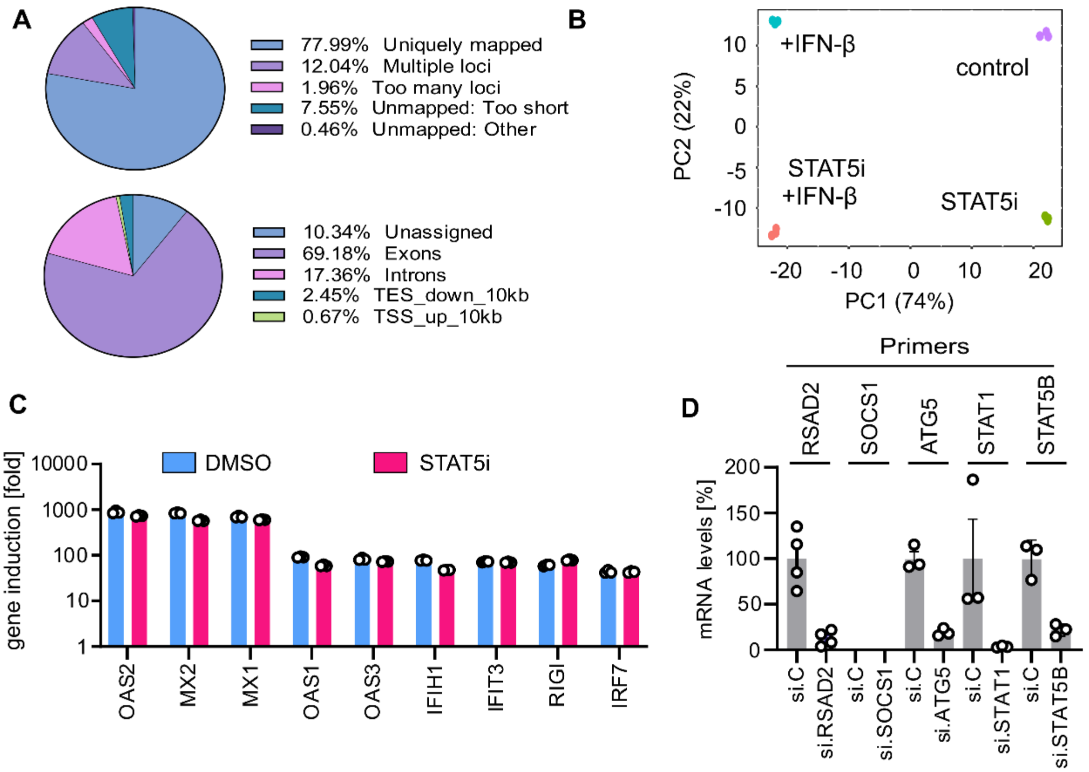


**Fig S5. Next-generation sequencing quality control, analysis and KD efficiency. A**, Percentage of mapped and unmapped reads for all samples n=12 (top panel). Percentage of read tags to genome features for all samples. n=12 (bottom panel). **B,** Principal Component Analysis of differential expression data, the individual samples are separated (violet control, cyan IFN-β, green STAT5 Inhibitor, orange IFN-β + STAT5 Inhibitor). n=3. **C,** Fold changes of selected ISGs from the data in Fig. 5A. n=3± SEM. **D,** RT-qRT-PCR analysis of siRNA-mediated knockdown of mRNA expression of data in Fig 5F. n=3-4± SEM.

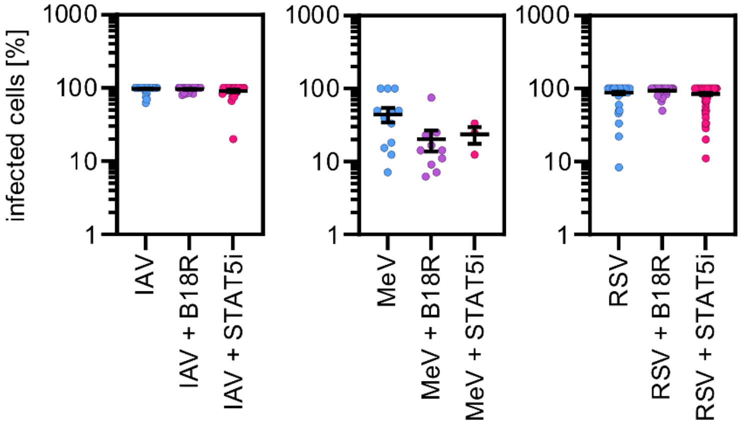


**Fig S6. Infection efficiencies**. Percentage of infected cells per tile of immunofluorescence data in Fig 6D. n=3-125± SEM.
